# CCRL2 promotes the interferon-γ signaling response in myeloid neoplasms with erythroid differentiation and mutated *TP53*

**DOI:** 10.1101/2025.02.21.639304

**Authors:** Nour Sabiha Naji, Sergiu Pasca, Theodora Chatzilygeroudi, Pablo Toledano-Sanz, Joseph Rimando, Yashvi Hemani, Brandy Perkins, Xinghan Zeng, Conover Talbot, Bogdan Paun, Abdulmuez Abdulmalik, Chen Lossos, Tatianna R Boronina, Ilias Sinanidis, Panagiotis Tsakiroglou, Priyanka Fernandes, Christopher Esteb, Alexander J. Ambinder, Robert N. Cole, Rena Xian, Ivana Gojo, Suman Paul, Mark J. Levis, Amy E. DeZern, Leo Luznik, Styliani Karanika, Linda S. Resar, Richard J. Jones, Frederick Bunz, Lukasz Gondek, Marios Arvanitis, Theodoros Karantanos

## Abstract

Patients with myeloid neoplasms with loss-of-function *TP53* mutations and erythroid differentiation have poor outcomes, and a better understanding of disease biology is required. Upregulation of interferon-γ (IFN-γ) signaling has been associated with acute myeloid leukemia (AML) progression and chemotherapy resistance, but its drivers remain unclear. In this study, we found that the surface receptor C-C motif chemokine receptor-like 2 (CCRL2) is overexpressed in AML with erythroid differentiation and *TP53* mutations compared to other AML subtypes and healthy hematopoietic cells. The knockout (KO) of CCRL2 suppressed erythroleukemia growth *in vitro* and *in vivo*. Further proteomics and transcriptomics analysis revealed IFN-γ signaling response as the top CCRL2-regulated pathway in erythroleukemia. Our mechanistic studies support direct CCRL2 driven IFN-γ signaling independent of exogenous IFN-γ, through phosphorylation of STAT1, via JAK2-dependent and independent mechanisms. CCRL2/IFN-γ signaling is upregulated in erythroid leukemias, and *TP53* mutated AML without concurrent increase of IFN-γ secretion in the bone marrow microenvironment and is directly induced by *TP53* KO. Finally, CCRL2/IFN-γ signaling is associated with the transformation of pre-leukemic single-hit *TP53* clones to multi-hit *TP53* mutated AML, increased resistance to venetoclax and worse survival in AML. Overall, our findings support that CCRL2 is an essential driver of cell-autonomous IFN-γ signaling response in myeloid neoplasms with erythroid differentiation and *TP53* mutations and highlight CCRL2 as a relevant novel target for these neoplasms.

**One Sentence Summary:** CCRL2 is overexpressed in AML with loss-of-function *TP53* mutations and erythroid differentiation and promotes IFN-γ signaling response via a cell-intrinsic mechanism.

## Introduction

Erythroid differentiation, bi-allelic *TP53* mutations, and loss of heterozygosity at the *TP53* gene are features associated with adverse biology in myeloid neoplasms[1-3]. Genomic characterization of erythroleukemia has revealed an exceptionally high prevalence of complex karyotype and bi-allelic *TP53* mutations[4, 5], suggesting a possible biologic link between loss of p53 function and erythroid differentiation in malignant hematopoiesis. Myeloid neoplasms with these features are characterized by poor response to chemotherapy and venetoclax-based therapies, high incidence of disease relapse, and very poor survival[1-3, 6, 7].

Recent studies have highlighted a possible implication of inflammatory signaling, including tumor necrosis factor-α (TNF-α) and interferon-γ (IFN-γ) in the induction of erythroid differentiation and evolution of *TP53* mutated acute myeloid leukemia (AML)[8, 9]. Consistently, upregulation of IFN-γ response signaling in AML blasts has been associated with worse survival and resistance to venetoclax[10]. However, the mechanism of inflammatory signaling upregulation in *TP53* mutated myeloid neoplasms remains unclear, with the central hypothesis being that it is mediated by dysregulation of the immune system in the bone marrow microenvironment[8].

C-C motif chemokine receptor-like 2 (CCRL2) is an atypical chemokine receptor involved in cell migration and inflammatory signaling activation, typically expressed on the surface of activated neutrophils and monocytes [11]. We found that CCRL2 is upregulated in the surface of CD34+ cells from patients with myelodysplastic syndrome (MDS) and blasts from patients with AML arising from MDS compared to healthy cells and de novo AML blasts[12]. Further, we have previously shown that *CCRL2* silencing with shRNA suppresses MDS/AML cell growth *in vitro* and *in vivo* and sensitizes them to hypomethylating agents [12, 13].

In this study, we show that AML with loss-of-function *TP53* mutations and erythroleukemias express the highest levels of CCRL2 across spectrum of AML subtypes, and that CCRL2 deletion by CRISPR-Cas9 suppresses their growth *in vitro* and *in vivo*. We also identified IFN-γ signaling as the top CCRL2-regulated pathway in these neoplasms, and demonstrated that *TP53* deletion directly induces CCRL2/IFN-γ upregulation without implication of the immune microenvironment, suggesting an endogenous activation of this pathway potentially mediated by intracellular regulators such as CCRL2 in *TP53* mutated MDS/AML cells.

## Results

### CCRL2 is upregulated in *TP53* mutated MDS/AML and erythroleukemia

The expression of CCRL2 in AML primary samples and cell lines was analyzed utilizing The Cancer Genome Atlas (TCGA)[14], Beat AML dataset [15]and DepMap Portal[16], which are publicly available datasets. Using TCGA data, we observed that French American British (FAB) M6 and M7 AML expressed higher levels of *CCRL2* compared to other AML subtypes (**Fig. 1A**). A comparison of the expression of *CCRL2* in AML samples (Beat AML dataset) showed that *TP53* mutated (MT) AMLs have higher *CCRL2* expression compared to wildtype (WT) ones (**Fig. 1B**). DepMap portal data analysis demonstrated that AML cell lines with erythromegakaryocytic differentiation (EM) have significantly higher CCRL2 expression than AML cell lines without EM differentiation (**Fig. 1C**).

**Fig. 1.**
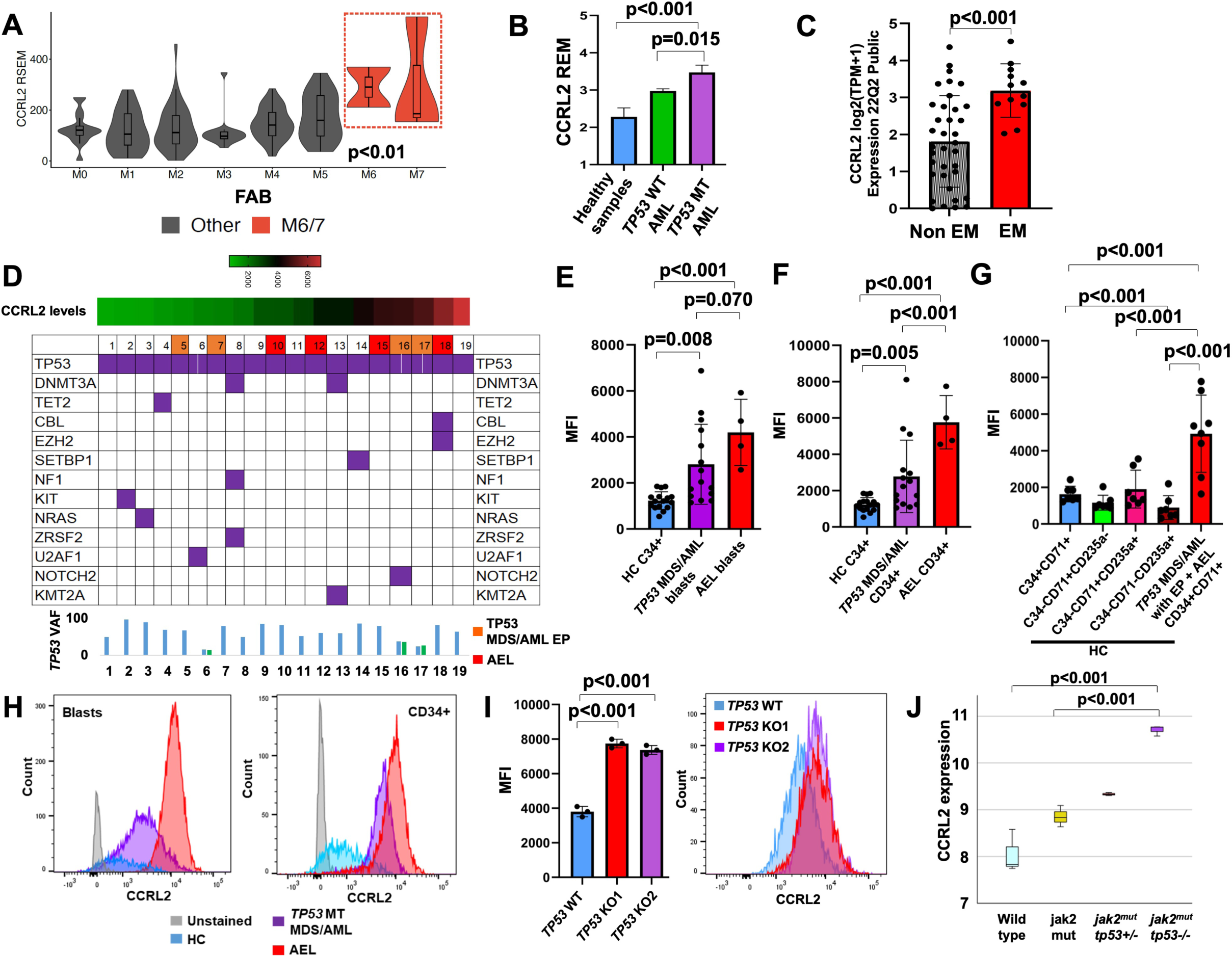
CCRL2 is upregulated in *TP53-*mutated MDS/AML and erythroleukemia. **(A)** Comparison of CCRL2 expression between different AML subtypes based on data extracted from the TCGA dataset. AML M6 and M7 subtypes, respectively, expressed higher levels of *CCRL2* compared to other AML subtypes based on RNA-Seq by Expectation-Maximization (RSEM) (p<0.01). **(B)** Comparison of the expression of *CCRL2* in AML samples based on Beat AML dataset showed that *TP53* mutated (MT) AMLs have higher *CCRL2* expression compared to wildtype (WT) ones (p=0.015) and healthy ones (p<0.001)**. (C)** Comparison of CCRL2 expression in different AML cell lines based on differentiation was performed using data from DepMap Portal. AML cell lines with EM differentiation have higher expression of *CCRL2* than AML cell lines with no EM differentiation(p<0.001). **(D)** Assessment of CCRL2 expression by flow cytometry in bone marrow samples of 15 patients with *TP53* MT MDS/AML with complex karyotype and either one *TP53* mutation of VAF at least 50% or two or more *TP53* mutations and 4 patients with AEL. 4 patients with *TP53* MT MDS/AML had erythroid predominance (EP) (≥50% of nucleated bone marrow cells were erythroid progenitors) but did not meet AEL criteria. **(E)** CCRL2 expression was compared in blasts from AEL patients (p<0.001) and *TP53* MT MDS/AML patients (p=0.008) to healthy controls **(F)** CD34+ cells from AEL patients (p<0.001) and *TP53* MT MDS/AML patients (p=0.005) have higher CCRL2 expression compared to healthy controls and CD34+ cells from AEL patients have higher CCRL2 expression than those from *TP53* MT MDS/AML patients(p<0.001). **(G)** CCRL2 expression in CD34+CD71+ cells from AEL patients and *TP53* MT MDS/AML patients with EP is significantly higher compared to erythroid progenitors from healthy donors (p<0.001). **(H)** Flow cytometry showing CCRL2 expression in blasts and CD34+ cells from representative samples of healthy donors, *TP53* MT MDS/AML, and AEL. **(I)** *TP53* knockout (KO) UKE-1 cells showed higher CCRL2 expression at the protein level compared to *TP53* WT UKE-1 cells (p<0.001). **(J)** Analysis of RNA-seq data (GSE180851) showed a gradual increase in CCRL2 expression in MEP cells associated with *Trp53* loss (from *Jak2*^mut^ to *Jak2*^mut^*Trp53*+/- and *Jak2*^mut^*Trp53*-/-) compared to wild type (WT) MEPs.

Next, we assessed CCRL2 expression in bone marrow samples of 15 patients with *TP53* MT MDS/AML with complex karyotype and either one *TP53* mutation of VAF at least 50% or two or more *TP53* mutations and 4 patients with AEL based on WHO 2022 criteria [17](**Fig. 1D**). Among the 15 *TP53* MT MDS/AML patients, 4 patients were identified to have erythroid predominance (EP) (≥50% of nucleated bone marrow cells were erythroid progenitors) but did not meet AEL criteria. Samples from 16 healthy donors were used as controls. Clinical, pathological, and molecular characteristics of healthy donors and patients are presented in **Fig. 1D**, **Supplementary Fig. 1A** and **Supplementary Tables 1 and 2**. The sorting strategy is shown in **Supplementary Fig. 1B, C**. CCRL2 expression was significantly higher in blasts and CD34+ cells from AEL patients and *TP53* MT MDS/AML patients than healthy controls (**Fig. 1E, F**). CCRL2 protein levels were higher in CD34+ cells from AEL patients than *TP53* MT MDS/AML patients (**Fig. 1F**). No significant correlation was found between CCRL2 expression and VAF or the presence of additional somatic mutations (**Fig. 1D**). *TP53* MT MDS/AML patients with EP showed a trend toward higher CCRL2 expression in their blasts and CD34+ cells compared to *TP53* MT MDS/AML without EP (**Supplementary Fig. 1D, E**). Finally, CD34+CD71+ cells from AEL patients and *TP53* MT MDS/AML patients with EP express significantly higher CCRL2 levels than healthy donors’ erythroid progenitors (**Fig. 1G**). Representative samples of healthy donors, *TP53* MT MDS/AML, and AEL are shown in **Fig. 1H**.

To further assess the effect of *TP53* deletion on CCRL2 expression in AML cells with erythroid differentiation, *TP53* was knocked out (KO) in UKE-1, an erythroleukemic cell line with WT *TP53*, by using CRISPR-Cas9 gene editing. By transducing UKE-1 cells with sgRNAs targeting *TP53* and selecting by treatment with blasticidin or nutlin-3a, two independent TP53 knockouts (KOs) were developed (KO1 and KO2) (**Supplementary Fig. 2A**). *TP53* KO UKE-1 cells demonstrated higher CCRL2 expression at protein (**Fig. 1I**) and RNA level (**Supplementary Fig. 2B**).

Finally, RNA-sequencing (RNA-seq) data from a published transgenic *TP53*-deleted erythroleukemia mouse model were analyzed to assess the impact of *TP53* deletion in CCRL2 expression. Li *et al.* recently demonstrated that deletion of *Trp53* (*Trp53*-/-) in Jak2^V617F^ knock-in (*Jak2*-mut) mice leads to the transformation of myeloproliferative neoplasm to AML with erythroid features derived from the megakaryocyte-erythroid progenitor (MEP) compartment[18]. Analysis of RNA-seq data (GSE180851) showed a gradual increase in CCRL2 expression in MEP cells associated with *Trp53* loss (from *Jak2*^mut^ to *Jak2*^mut^*Trp53*+/- and *Jak2*^mut^*Trp53*-/-) compared to WT MEPs **(Fig. 1J**).

### CCRL2 increases the clonogenicity of erythroleukemia cells *in vitro* and promotes their growth *in vivo*

To assess the effects of CCRL2 on the clonogenicity of erythroleukemia cells, CCRL2 was knocked out in two erythroleukemic cell lines, TF-1 and F36P, one BCR-ABL+ AML cell line with erythroid features, K562 and two *JAK2^V617F^* mutated erythroleukemic cell lines, SET2 and HEL, by using two sgRNAs targeting CCRL2 (sgCCRL2 1 and sgCCRL2 2) and scrambled sgRNA (sgControl) as control. Suppression of CCRL2 expression was confirmed by flow cytometry (**Supplementary Fig. 2B-G**). CCRL2 KO suppressed the clonogenicity of TF-1, F36P, K562, SET2 and HEL cells (**Fig. 2A-E**). Conversely, CCRL2 KO showed no effect on the clonogenicity of MV4-11 cells, which are *TP53* WT cells with monocytic differentiation and express low levels of CCRL2 (**Supplementary Fig. 2H-I**).

**Fig. 2.**
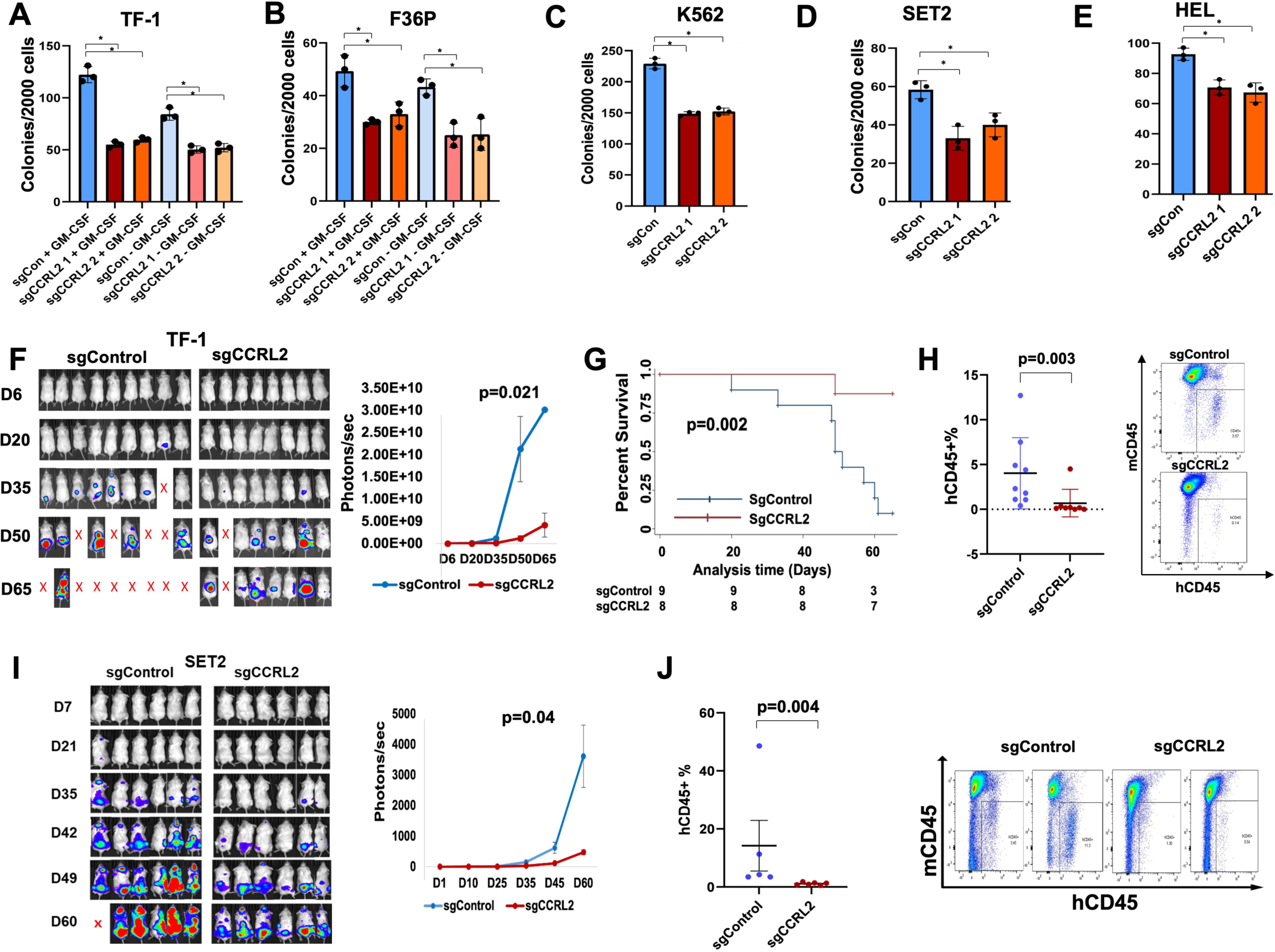
CCRL2 promotes the clonogenicity of erythroleukemia cells *in vitro* and the growth of erythroleukemia in mice. **(A-B)** CCRL2 KO with two different sgRNAs decreases the colony formation of the GM-CSF dependent TF-1 and F36P cells in the absence and presence of granulocyte-macrophage colony-stimulating factor (GM-CSF). **(C-E)** CCRL2 KO with two different sgRNAs decreases the colony formation of K562, SET2, and HEL cells. **(F)** TF-1 cells with WT (sgControl) or KO (sgCCRL2) CCRL2 were transduced with a GFP+/Luciferase dual reporter retrovirus and injected into NSG mice. At day 50 after injection, the bioluminescence signal in mice injected with CCRL2 KO was significantly lower (p=0.021). **(G)** Mice engrafted with sgControl cells had a significantly poorer overall survival than those engrafted with sgCCRL2 (p=0.002). **(H)** On day 65, survived mice were euthanized. Disease burden in bone marrow was measured by human CD45+%. Mice engrafted with sgCCRL2 had a significantly lower percentage of human CD45+ than those engrafted with sgControl (p=0.003). **(I)** SET2 cells with WT (sgControl) or KO (sgCCRL2) CCRL2 were transduced with a GFP+/Luciferase dual reporter retrovirus and injected into NSG mice. At day 60 after injection, the bioluminescence signal in mice injected with suppressed CCRL2 was significantly lower (p=0.04). **(J)** Disease burden in bone marrow was measured by human CD45+%. Mice engrafted with sgCCRL2 had a significantly lower percentage of human CD45+ than those engrafted with sgControl (p=0.004).

To evaluate the effect of CCRL2 KO in the growth of erythroleukemic cells *in vivo*, GFP/Luciferase+ TF-1 cells transduced with sgControl or sgCCRL2 (sgCCRL2 1) were injected intravenously in NOD.Cg-*Prkdc^scid^ Il2rg^tm1Wjl^*/SzJ (NSG) mice. No significant differences in engraftment rate were observed, but leukemic growth of sgCCRL2 cells was significantly suppressed (**Fig. 2F**), and those mice had a significantly improved survival rate (**Fig. 2G**). On day 65, surviving mice were euthanized, and sgCCRL2 mice exhibited a significantly lower disease burden in their bone marrows measured by human CD45+% (**Fig. 2H**). Similarly, GFP/Luciferase+ SET2 transduced with sgControl or sgCCRL2 were engrafted in NSG mice. The CCRL2 KO group exhibited decreased leukemic growth, as measured by the bioluminescence signal (**Fig. 2I**). Disease burden measured by the percentage of human CD45+ in the bone marrow was decreased in mice engrafted with SET2 KO cells (**Fig. 2J**). Overall, these findings suggest that CCRL2 induces the growth and clonogenicity of leukemic cells with erythroid differentiation both *in vitro* and in NSG mice.

### CCRL2 promotes IFN-γ response signaling in erythroleukemia cells

To assess the most prominent effect of CCRL2 KO on erythroleukemia cells at the protein level, unbiased phospho-proteomics analysis was performed in TF-1 cells transduced with either sgControl or one of two independent sgRNAs (two replicates (KO1 and KO2) of sgCCRL2 1 and one replicate (KO3) of sgCCRL2 2).

Pathway enrichment analysis of the phospho-proteomics data identified IFN signaling was identified as the top CCRL2-regulated pathway, a pathway sharing various molecules with hyperchemokinemia in influenza, the second most prominent pathway (**Fig. 3A**). Liver X Receptor-Retinoid X Receptor (LXR/RXR) was the top pathway suppressed by CCRL2 expression. The IFN signaling regulator STAT1 (both long and short isoforms) along with the IFN-γ targets IFIT1, IFIT3, and ISG15 were amongst the top CCRL2 targets, while various LXR/RXR targets were upregulated by CCRL2 KO (**Fig. 3B**). A negative interaction between IFN-γ and LXR/RXR pathways has been described[19, 20], providing some rationale for our finding that CCRL2 promotes IFN signaling and suppresses LXR/RXR (**Supplementary Fig. 3A**). Besides these top targets, various IFN-γ/STAT1 targets are significantly downregulated by CCRL2 KO (**Fig. 3C**).

**Fig. 3.**
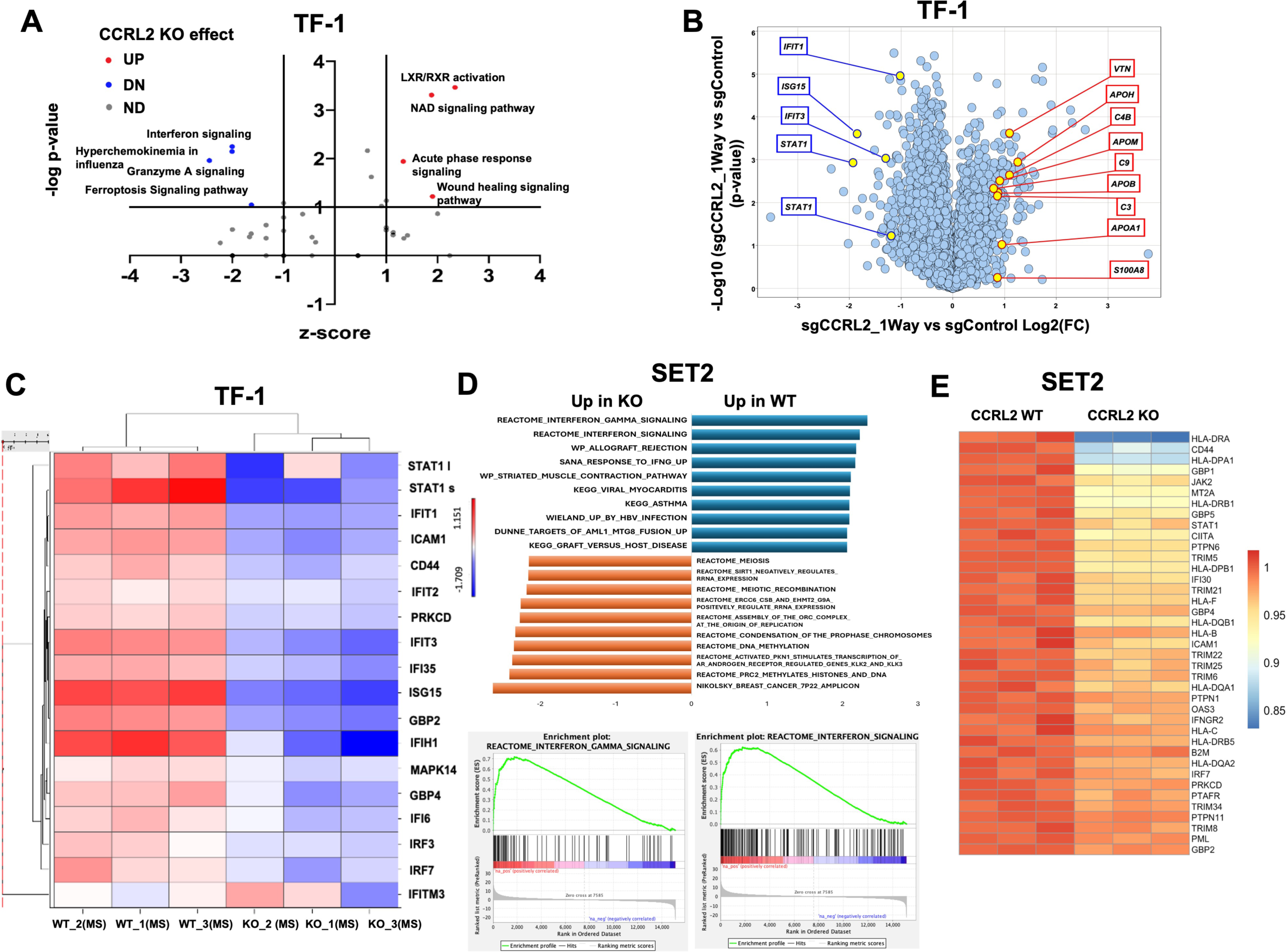
CCRL2 promotes IFN-γ response signaling in erythroleukemia cells. **(A)** Pathway enrichment analysis of phospho-proteomic data was performed to identify the pathways affected more prominently by CCRL2 KO at the protein level in TF-1 cells. IFN signaling was identified as the top pathway downregulated by CCRL2 KO, and Liver X Receptor-Retinoid X Receptor (LXR/RXR) pathway was the top pathway upregulated by CCRL2 KO. **(B)** The IFN-γ signaling regulator STAT1 (both long and short isoforms) along with the IFN-γ targets (IFIT1, IFIT3 and ISG15) were amongst the top downregulated proteins by CCRL2 KO. LXR/RXR targets are upregulated by CCRL2 knockout (KO). **(C)** Heatmap, including the top CCRL2 regulated targets represented in the volcano plot in addition to other various IFN-γ/STAT1 targets, shows that these targets are significantly downregulated by TF-1 CCRL2 KO. MS represents mean subtracted values. The intensity scale to the right shows a range of values from -1.709 to 1.151, with the blue color representing the lower extreme values of this range and red color representing the high extreme values of the range. **(D)** IFN-γ signaling was found to be the top CCRL2-regulated pathway by gene set enrichment analysis of RNA-seq data in SET2 cells transduced with sgControl or sgCCRL2 (CCRL2 KO). **(E)** Heatmap shows that CCRL2 regulated IFN-γ/STAT1 targets, are significantly downregulated by SET2 CCRL2 KO. The intensity scale to the right shows a range of values from 0.85 to 1, with the blue color representing the lower extreme values of this range and the red color representing the high extreme values of the range.

Next, bulk RNA-seq was performed followed by gene set enrichment analysis (GSEA) using a compilation of pathways from MSigDB[21] in CCRL2 WT or KO SET2 cells (**Supplementary Fig. 3B, C**). IFN-γ gene network activation was the top CCRL2-regulated pathway (**Fig. 3D-E, Supplementary Fig. 3D**). These results further support that IFN-γ response signaling is the top CCRL2-regulated pathway in erythroleukemia at both RNA and protein levels.

To validate our findings, the phosphorylation of STAT1, the main IFN-γ signaling regulator[22] in Tyrosine 701 (Y701) and Serine 727 (S727) was measured in CCRL2 WT and KO TF-1, F36P, and SET2 cells showing significant suppression of STAT1 phosphorylation by CCRL2 KO (**Fig. 4 A-C**). Of note, S727 STAT1 phosphorylation was not detected in F36P cells, and Y701 phosphorylation was detected in these cells only in the presence of GM-CSF (**Fig. 4B**). CCRL2 KO in TF-1 cells suppressed STAT1 nuclear translocation in TF-1 cells (**Supplementary Fig. 4A**). Consistently, quantitative real-time PCR showed that CCRL2 KO suppresses the RNA levels of 3 representative IFN-γ/STAT1 target genes, namely *IFIT3*, *IFI30* and *MT2A* (**Fig. 4D**).

**Fig. 4.**
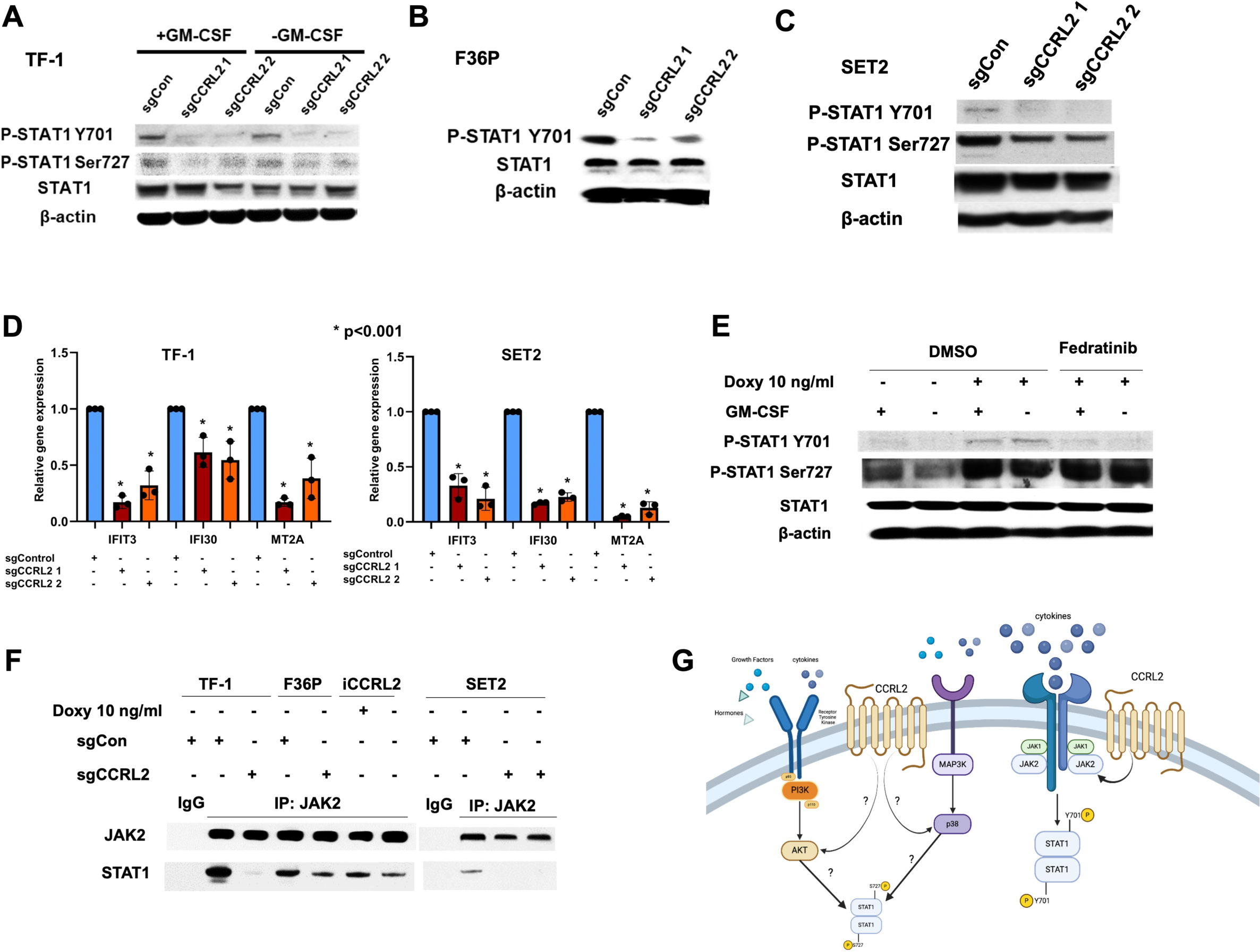
CCRL2 induces STAT1 phosphorylation via JAK2-dependent and independent mechanisms. **(A)** Western blotting was performed to analyze two sites of STAT1 phosphorylation: tyrosine 701 (Y701) phosphorylation and serine 727 (S727) phosphorylation. CCRL2 KO suppressed STAT1 Y701 phosphorylation in TF-1 and had a less prominent effect on STAT1 S727 phosphorylation. **(B)** Similarly, CCRL2 KO suppressed STAT1 Y701 phosphorylation in F36P cells. No S727 phosphorylation was detected in F36P cells, and no STAT1 phosphorylation was detected in the absence of GM-CSF. **(C)** STAT1 Y701 and S727 phosphorylation was suppressed in SET2 CCRL2 KO cells. **(D)** Quantitative real-time PCR demonstrated that CCRL2 KO suppresses the RNA levels of 3 representative IFN-γ/STAT1 target genes, namely *IFIT3*, *IFI30* and *MT2A* in TF-1 and SET2 cells compared to WT cells. **(E)** Treatment of iCCRL2 TF-1 cells with 10 ng/ml doxycycline induced STAT1 Y701 and S727 phosphorylation; however, concurrent treatment with fedratinib suppressed Y701 but not S727 phosphorylation. **(F)** Co-immunoprecipitation assay showing that CCRL2 KO decreases the precipitation of STAT1 with an anti-JAK2 antibody in TF-1, F36P cells, and SET2 while induction of CCRL2 by 10 ng/ml doxycycline increases STAT1 precipitation in iCCRL2 TF-1 cells. **(G)** CCRL2 induces STAT1 Y701 phosphorylation via JAK2, potentially promoting STAT1 S727 phosphorylation via JAK2-independent pathways.

Given that exogenous IFN-γ is critical for the activation of IFN-γ signaling response in the bone marrow microenvironment[10], the effect of CCRL2 on the sensitivity of AML cells to exogenous IFN-γ was assessed. CCRL2 KO did not affect the upregulation of *IFIT3* expression in TF-1 cells as a response to exogenous IFN-γ (**Supplementary Fig. 4B**). Thus, our findings suggest that CCRL2 promotes a cell intrinsic activation of IFN-γ response signaling rather than the sensitivity of leukemic cells to exogenous IFN-γ.

Our group has previously shown that CCRL2 promotes JAK2/STAT signaling[12]. The regulation of IFN-γ signaling at the molecular level is complex, involving numerous pathways apart from JAK2/STAT[23]. Mainly, Y701 phosphorylation of STAT1 is mediated by JAK1/2, but S727 phosphorylation is promoted by AKT and p38/MAPK pathways[24-26]. To investigate possible mediators of the CCRL2-induced STAT1 phosphorylation, our doxycycline-inducible CCRL2 TF-1 cells model [13] was used. Particularly, iCCRL2 TF-1 cells were treated with 0 or 10 ng/ml doxycycline to induce CCRL2 expression, leading to the induction of cell growth (**Supplementary Fig. 4C**). Treatment of iCCRL2 TF-1 cells with 10 ng/ml doxycycline induced STAT1 Y701 and S727 phosphorylation, while concurrent treatment with the JAK2 inhibitor fedratinib suppressed Y701 but not S727 phosphorylation (**Fig. 4E**). This finding suggests that CCRL2 promotes Y701 STAT1 phosphorylation at least partially via JAK2 activation but probably promotes S727 STAT1 phosphorylation via other JAK2-independent pathways. Consistently with our previous findings[12], co-immunoprecipitation showed that CCRL2 KO suppressed JAK2 and STAT1 interaction in TF-1, F36P, and SET2 cells, while doxycycline treatment induced JAK2/STAT1 interaction in doxycycline-inducible CCRL2 TF-1 cells (**Fig. 4E**).

We have previously shown that CCRL2 activates STAT3/5 via direct binding to JAK2 and induction of JAK2/STAT interaction[12]. Co-immunoprecipitation using an anti-JAK2 antibody revealed that CCRL2 KO in TF-1, F36P, and SET2 cells suppresses the interaction between JAK2 and STAT1 while induction of CCRL2 expression increases the interaction between these two molecules in TF-1 cells (**Fig. 4F**).

Overall, these findings support that IFN-γ signaling is the top CCRL2-regulated pathway in erythroleukemia cells and that CCRL2 activates a cell-intrinsic IFN-γ response signaling without affecting the response of leukemic cells to exogenous IFN-γ by promoting STAT1 phosphorylation partially via JAK2 but also via JAK2-independent pathways (**Fig. 4G**).

### IFN-*γ* response signaling is upregulated in AML with erythroid differentiation and *TP53* mutations but is not associated with increased IFN-γ secretion in the bone marrow microenvironment

It was recently reported, that IFN-γ response signaling is upregulated in monocytic AML and AML with deletions in chromosome 7. However, although IFN signaling is upregulated in TP53 mutation, its specific association with erythroid AML carrying TP53 mutations has not been described.

To evaluate the level of IFN-γ response signaling activation in AML with erythroid features and *TP53* mutations, an overall score for 18 genes commonly implicated in IFN-γ signaling and identified as CCRL2 targets by our analysis (*CCRL2, STAT1, IFIT1, ICAM1, CD44, IFIT2, PRKCD, IFIT3, IFI35, ISG15, GBP2, IFIH1, MAPK14, GBP4, IFI6, IRF3, IRF7, IFITM3*) was calculated. Based on TCGA data, AML with erythroid and megakaryocytic (EM) differentiation, FAB M6 and M7, respectively, had a higher overall score than other subtypes of AML (**Fig. 5A**). Similarly, data analysis from DepMap Portal showed higher levels of CCRL2/IFN-γ signaling score in AML cell lines with erythroid differentiation than in other AML cell lines (**Fig. 5B**). To assess the expression of CCRL2/IFN-γ targets in *TP53* mutated AML, we used the Beat AML dataset to compare CCRL2/IFN-γ signaling overall score between *TP53* MT and WT AML*. TP53* MT AML had a higher CCRL2/IFN-γ signaling score than *TP53* WT AML (**Fig. 5C**). CCRL2/IFN-γ signaling score was also higher in *TP53* WT AML samples than healthy CD34+ cells (**Fig. 5C**). Among the top CCRL2-regulated genes *IFIT1*, *IFIT3,* and *ISG15*, *IFIT1* was also found to be significantly overexpressed in *TP53* MT AML samples compared to *TP53* WT ones, and a similar trend was observed for *IFIT3* and *ISG15* (**Supplementary Fig. 5A**). Of note, all three genes were upregulated in AML samples compared to healthy bone marrow mononuclear cells (**Supplementary Fig. 5A**).

**Fig. 5.**
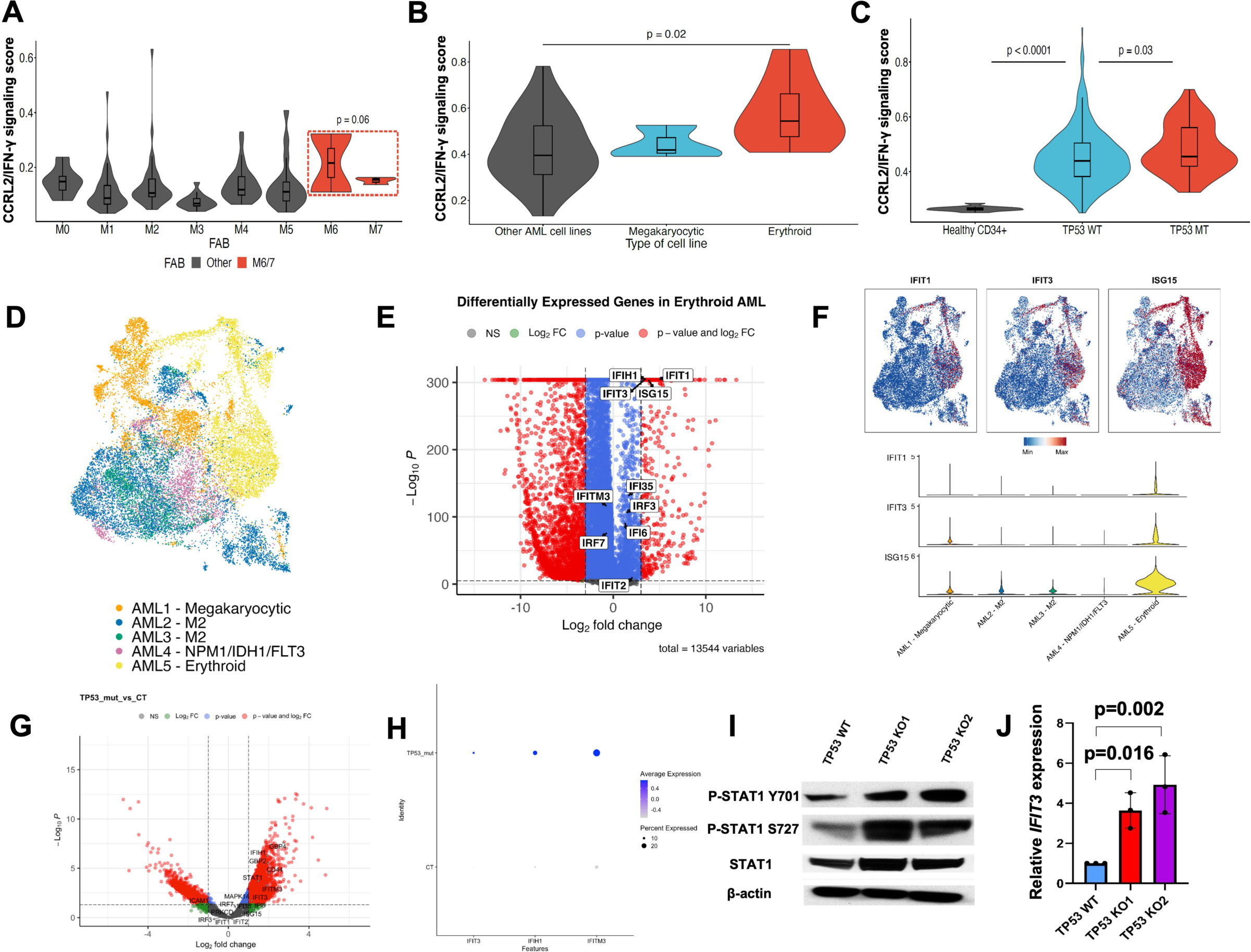
CCRL2/IFN-γ signaling is upregulated in AML with erythroid differentiation and *TP53* mutations without an increase in IFN-γ secretion. **(A)** Based on data derived from TCGA, an overall score for 18 genes involved in CCRL2/IFN-γ signaling was calculated. AML with erythroid and megakaryocytic (EM) differentiation (AML M6 and M7 subtypes, respectively) expressed higher levels of the CCRL2/IFN signaling score (p=0.06) compared to other subtypes of AML. **(B)** Analysis of DepMap Portal data showed that erythroid cell lines had higher CCRL2/IFN-γ signaling score than other AML cell lines (p=0.02). (**C)** Analysis of Beat AML data demonstrated that AML with *TP53* mutation (MT) had a higher CCRL2/IFN-γ signaling score compared to *TP53* wildtype (WT) AML (p=0.03). CCRL2/IFN-γ signaling score was also higher in *TP53* WT AML than healthy CD34+ controls(p<0.0001). **(D)** Based on publicly available single-cell RNA sequencing data, clustering of AML blasts and monocytes from 5 patients with different AML subtypes was performed based on differentiation. **(E)** Analysis of single-cell RNA sequencing data showed that blasts with erythroid differentiation express higher levels of various CCRL2/IFN-γ targets, including *IFIT1*, *IFIT3,* and *ISG15,* compared to other AML differentiation clusters. **(F**) The AML patient with erythroid differentiation had upregulated expression of *IFIT1* (average log2 fold change = 4.98; adjusted p < 0.0001); *ISG15* (average log2 fold change = 3.80; adjusted p < 0.0001); and *IFIT3* (avg log2 fold change = 3.37; adjusted p < 0.0001) compared to the other AML patients. **(G)** Based on analysis of another publicly available single-cell RNA sequencing dataset, the volcano plot shows that the expression of most CCRL2/IFN-γ targets is higher in *TP53* mutated AML than *TP53* WT AML**. (H)** Specifically, *IFIT3, IFIH1,* and *IFITM3* are significantly upregulated in blasts from *TP53* mutated AML samples compared to blasts from *TP53* WT AML patients. **(I)** *TP53* KO in UKE-1 cells led to a prominent increase in both Y701 and S727 phosphorylation of STAT1 compared to *TP53* WT UKE-1 cells. **(J)** *IFIT3* is upregulated in *TP53* KO UKE1 cells compared to *TP53* WT UKE-1 cells (p=0.016 with KO1 and p=0.002 with KO2).

To assess the level of CCRL2/IFN-γ signaling activation in primary samples, publicly available single-cell RNA-seq (scRNAseq) datasets were analyzed. First, an analysis of a publicly available scRNAseq dataset of AML blasts and monocytes from five AML patients that was previously published[27] was performed. Two patients in the dataset had megakaryocytic (AML1) and erythroid (AML5) AML; the other three patients (AML2-4) had non-megakaryocytic, non-erythroid AML (**Fig. 5D**). Most of the CCRL2/IFN-γ targets were found to be expressed at a relatively higher level in blasts with erythroid differentiation (**Fig. 5E, Supplementary Fig. 5B**), with the top CCRL2/IFN-γ targets *IFIT1*, *IFIT3*, and *ISG15* being among the significantly different ones (**Fig. 5E**). Compared with the other four patients, the AML patient with erythroid differentiation had upregulated expression of *IFIT1* (average log2 fold change = 4.98; adjusted p < 0.0001); *ISG15* (average log2 fold change = 3.80; adjusted p < 0.0001); and *IFIT3* (avg log2 fold change = 3.37; adjusted p < 0.0001) (**Fig. 5F**).

Next, we reanalyzed scRNAseq data from a study by van Galen *et al.* [28] who performed scRNA-seq in bone marrow aspirates of 16 AML patients including 3 individuals with *TP53* mutations. Blasts were identified by CD34 and c-KIT (CD117) expression (**Supplementary Fig. 5C**). Most of the CCRL2/IFN-γ targets were overexpressed in *TP53* mutated AML cells compared to *TP53* WT ones (**Fig. 5G**), with *IFIT3*, *IFIH1* and *IFITM3* being the ones with the most notable differences (**Fig. 5H**). Using the same dataset, the expression of *IFN*-γ in CD3+ cells (T-cells) (**Supplementary Fig. 5C**) was compared between *TP53* WT and MT AML patients, showing that patients with *TP53* MT AML expressed relatively lower levels of *IFN-* γ compared to the WT ones without difference reaching statistical significance (**Supplementary Fig. 5D**).

To further assess if induced IFN-γ response signaling in *TP53* mutated AML is associated with increased IFN-γ secretion in the bone marrow microenvironment, CD3+CD45+ and CD3-CD45+ cells were sorted from 4 *TP53* mutated AML patients and 3 healthy bone marrow donors (**Supplementary Table 1, Supplementary Fig. 6A**). Blasts by dim CD45 and low side scatter were also sorted from the *TP53* mutated AML patients (**Supplementary Fig. 6A**). Following 72 hours of cell culture (50,000 cells/ml), the levels of IFN-γ were measured by ELISA, showing that T-cells from *TP53* MT AML cells secreted relatively lower IFN-γ levels compared to healthy donors (**Supplementary Fig. 6B**).

Since a cell-intrinsic event potentially links *TP53* mutations with upregulation of CCRL2/IFN-γ signaling, activation of STAT1 phosphorylation and *IFIT3* RNA levels were compared between *TP53* WT and KO UKE1 cells. *TP53* KO led to a prominent increase in both Y701 and S727 phosphorylation of STAT1 (**Fig. 5I**) and upregulation of the IFN-γ target *IFIT3* (**Fig. 5J**) in UKE1 cells. These results are consistent with the observed increase in the expression of CCRL2 in UKE1 cells following *TP53* KO (**Fig. 1I**).

Overall, these findings support that CCRL2/IFN-γ targets are upregulated in AML cells with erythroid differentiation and *TP53* mutations compared to other AML subtypes without increased secretion of IFN-γ in the bone marrow microenvironment, suggesting a possible cell-intrinsic mechanism involved in the upregulation of this pathway. Particularly, *TP53* deletion leads directly to the induction of STAT1 phosphorylation and upregulation of CCRL2/IFN-γ targets.

### CCRL2/IFN-γ signaling upregulation is associated with selection of *TP53* mutated clones and resistance to venetoclax in AML

Given that the role of IFN-γ response signaling in cancer progression remains unclear[29, 30], we then asked what the possible functional implication of CCRL2/IFN-γ signaling response activation in *TP53* mutated AML is.

First, analysis of DepMap portal publicly available CRISPR-Cas9 dataset [16]revealed that the majority of CCRL2/IFN-γ target genes demonstrate a negative gene effect in AML cell lines with erythroid differentiation and *TP53* mutations (HEL, HEL9217, SET2, TF-1, CMK115, MUTZ8, OCIM2, and F36P) (**Fig. 6A**). On the contrary, treatment of primary AML blasts sorted from 3 independent patients with TP53 mutated AML (**Supplementary Table 1**) with exogenous IFN-γ for 72 hours induced apoptosis of leukemic blasts and decreased the percentage of CD34+CD117+ cells (**Fig. 6B**). These findings further support that there may be a functional distinction between cell intrinsic upregulation of IFN-γ signaling and the response of leukemic cells to exogenous IFN-γ in the bone marrow microenvironment.

**Fig. 6.**
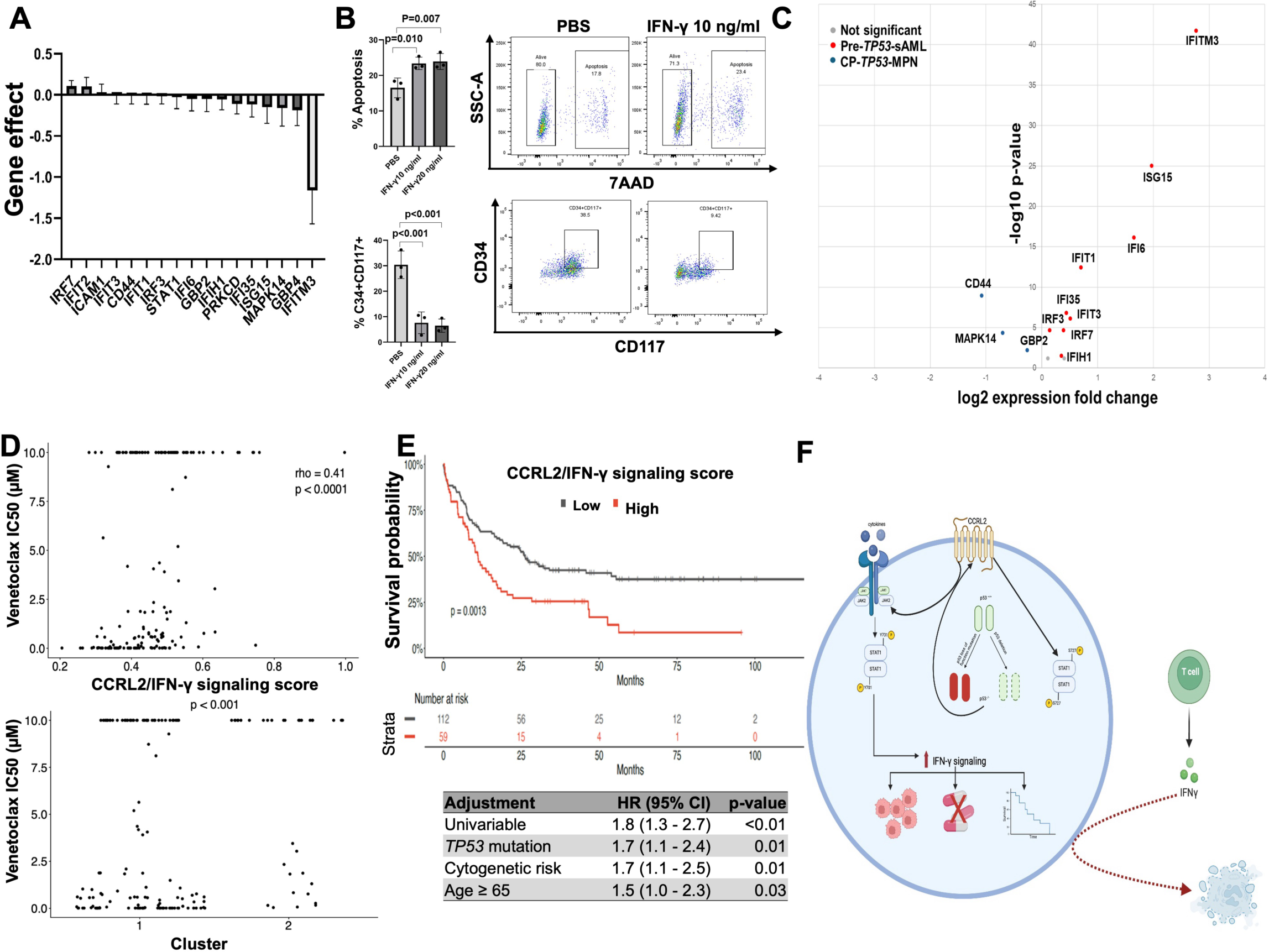
CCRL2/IFN-γ signaling upregulation is associated with selection of *TP53* mutated clones and AML resistance to venetoclax. **(A)** Analysis of DepMap portal publicly available CRISPR-Cas9 dataset showed that most CCRL2/IFN-γ target genes exhibit a negative gene effect in AML cell lines with erythroid differentiation and *TP53* mutations (HEL, HEL9217, SET2, TF-1, CMK115, MUTZ8, OCIM2 and F36P) (**B)** Treatment of primary AML blasts from 3 independent *TP53* mutated AML patients with 10 ng/ml and 20 ng/ml of IFN-γ for 72 hours induced apoptosis of leukemic blasts (p=0.010 and p=0.007 respectively) and decreased the percentage of CD34+CD117+ cells (p<0.001 for both doses). **(C)** Volcano plot showed that a significant subset of CCRL2/IFN-γ targets is upregulated in pre-leukemic *TP53* heterozygous clones from MPN patients who transformed to sAML compared to those who remained in CP-MPN. **(D)** Analysis of data from Beat AML dataset showed that CCRL2/IFN-γ signaling score is positively correlated with venetoclax IC50 (p<0.0001). Patients in cluster 2 who had a high expression of *IFIT2, IFIT1, IFIT3, STAT1, IFI6, IFIH1, ISG15,* and *IRF7* had significantly higher Venetoclax IC50 values compared to cluster 1 patients (p<0.001). **(E)** Based on data from Beat AML dataset, patients with high CCRL2/IFN-γ signaling score (red) had a worse overall survival compared to patients with low CCRL2/IFN-γ signaling score (gray) over 100 months. **(F)** *TP53* deletion/loss of function promotes CCRL2/IFN-γ signaling response in AML subtypes, which appears to be associated with a cell-intrinsic effect independent of IFN-γ levels in the microenvironment, and the latter is associated with selection of *TP53* mutated pre-leukemic clones, resistance to venetoclax and poor overall survival.

It was recently reported that inflammatory signaling might contribute to preleukemic clonal evolution toward *TP53* multi-hit mutated sAML with erythroid features[8]. Analysis of scRNA-seq data from this report showed that pre-leukemic single-hit *TP53* heterozygous clones from patients with myeloproliferative neoplasms (MPN) that transformed to multi-hit *TP53* mutated sAML have higher expression of the top CCRL2/IFN-γ targets *IFIT1*, *IFIT3* and *ISG15* compared to *TP53* heterozygous clones from patients who remained in chronic phase (CP-MPN) (**Supplementary Fig. 6C**). Similarly, a significant subset of CCRL2/IFN-γ targets were upregulated in pre-leukemic single-hit *TP53* heterozygous clones from MPN patients that transformed to sAML compared to those who remained in CP-MPN (**Fig. 6C**). These results support that CCRL2/IFN-γ signaling may be implicated in the progression of AML with erythroid features via selection of *TP53* mutated pre-leukemic clones.

Wang *et al.* recently showed that overexpression of IFN-γ targets and particularly *IFITM3* is associated with lower sensitivity to venetoclax [10]. Other studies have highlighted that *TP53* mutations and erythroid differentiation are associated with higher rates of venetoclax resistance[27]. Analysis of data from Beat AML dataset showed that CCRL2/IFN-γ signaling score is positively correlated with venetoclax IC50, supporting that upregulation of this pathway is associated with increased resistance to venetoclax (**Fig. 6D**). Based on the expression pattern of the CCRL2/IFN-γ targets. Patients were separated into two clusters; cluster 2 included those with a high expression of *IFIT2, IFIT1, IFIT3, STAT1, IFI6, IFIH1, ISG15, IRF7* while cluster 1 included the rest of the patients (**Supplementary Fig. 6D**). Patients in cluster 2 had significantly higher venetoclax IC50 values compared to cluster 1 patients (**Fig. 6D**), suggesting that the most direct CCRL2/IFN-γ targets, including *IFIT1*, *IFIT3,* and *ISG15* are potentially more potent mediators of venetoclax resistance.

Finally, the impact of CCRL2/IFN-γ signaling score on the survival of AML patients was assessed using data from Beat AML, demonstrating a negative impact of this score on the overall survival of AML patients (**Fig. 6E**). Multivariable analysis showed that this effect is independent of TP53 mutation, cytogenetic risk, and patients’ age (**Fig. 6E**).

Taken together, our data support that CCRL2 is overexpressed in AML with erythroid differentiation and *TP53* mutations and induces leukemia growth. CCRL2 promotes IFN-γ signaling response in these AML subtypes, which appears to be associated with a cell-intrinsic effect that is independent of IFN-γ levels in the microenvironment. CCRL2/IFN-γ targets are upregulated in erythroid and *TP53*-mutated AML, and this signaling is associated with the selection of *TP53*-mutated pre-leukemic clones, resistance to venetoclax and poor overall survival (**Fig. 6F**).

## Discussion

Patients with myeloid neoplasms with erythroid differentiation and *TP53* mutations exhibit very short survival due to poor response to therapies, including venetoclax and high incidence of relapse [3, 31-37]. Therefore, a better understanding of disease pathogenesis is required. The role of IFN-γ signaling in myeloid neoplasms remains unclear, but it was recently reported that inflammatory pathways may be implicated in the selection of *TP53* mutated pre-leukemic clones driving disease progression[8]. Upregulation of IFN-γ signaling was also associated with AML progression and resistance to venetoclax[10]. However, the mechanisms implicated in the regulation of IFN-γ signaling in myeloid neoplasms remain unclear, and the leading hypothesis is that secreted IFN-γ in the marrow microenvironment is the primary driver of this pathway.

Here, we found that the surface receptor CCRL2 is overexpressed in MDS/AML with erythroid differentiation and *TP53* mutations associated with complex karyotype compared to other AML subtypes and healthy hematopoietic cells, and its deletion suppresses erythroleukemic cells clonogenicity *in vitro* and growth *in vivo*. CCRL2 is normally expressed in activated macrophages, neutrophils, and NK cells and promotes inflammatory signaling [11, 38, 39]. We previously found that CCRL2 is overexpressed in progenitors from MDS patients and sAML blasts [12]. Erythroleukemia shares methylation and transcriptomic features with both AML and MDS [32], suggesting a possible overlap of erythroleukemia biology with MDS and sAML, consistent with our findings. Recent data highlight that the clinical outcomes of MDS/AML spectrum are primarily affected by specific biological characteristics, with loss of p53 function having the most prominent effects and not by the exact blasts’ percentage[40]. Based on our published data [12]and our current results, CCRL2 overexpression is probably associated with adverse disease biology within this spectrum, supporting a correlation with loss of p53 function.

Phospho-proteomics and transcriptomics analysis highlighted IFN-γ response signaling as the top CCRL2-regulated pathway in erythroleukemia cells. Activation of inflammatory pathways is consistent with the well-described role of CCRL2 in promoting inflammatory response in healthy differentiated cells [11, 38, 39]. Our studies confirmed suppressed nuclear translocation of STAT1 and decreased STAT1 target expression by CCRL2 KO in erythroleukemia cells. Mechanistically, JAK2 is a critical upstream regulator of STAT1 phosphorylation and IFN-γ signaling [41], and we have reported that CCRL2 induces JAK2/STAT signaling[12]. Here, by utilizing our doxycycline-inducible CCRL2 TF-1 cells, we found that CCRL2 induces STAT1 Y701 phosphorylation via JAK2 but also promotes STAT1 S727 phosphorylation via a JAK2 independent mechanism. While Y701 phosphorylation of STAT1 is mediated by JAK2, other pathways, including p38/MAPK and AKT, induce S727 phosphorylation[24, 25]. Further studies are required to investigate a possible implication of CCRL2 in regulating these pathways mediating STAT1 S727 phosphorylation. Of note, we found that CCRL2 promotes IFN-γ response signaling in erythroleukemic cells but does not affect their response to exogenous IFN-γ, suggesting a possible distinction between cell-intrinsic and extrinsic regulation of IFN-γ signaling.

Our developed CCRL2/IFN-γ gene set was upregulated in AML with erythroid differentiation and *TP53* mutations based on TCGA, DepMap portal, and Beat AML data. By analyzing scRNAseq data from publicly available datasets, we showed increased expression of CCRL2/IFN-γ targets in blasts from erythroid leukemia and *TP53* MT AML compared to other AML subtypes. Importantly, we did not observe any increase of *IFNG* expression in T-cells from *TP53* MT AML samples compared to *TP53* WT samples, and sorted T-cells from *TP53* MT AML patients secreted relatively lower levels of IFN-γ compared to healthy donors. This result is consistent with previous studies reporting lower secretion of IFN-γ by T-cells [42]and lower IFN-γ/IFN-γ receptor interactions in *TP53* MT AML samples[10]. These results further support a possible cell-intrinsic mechanism of IFN-γ response signaling upregulation in *TP53* MT myeloid neoplasms. Indeed, we demonstrated that *TP53* KO causes an increase in CCRL2 levels, STAT1 phosphorylation, and *IFIT3* expression, suggesting that loss of p53 can promote CCRL2/IFN-γ signaling activation through a cell-intrinsic phenomenon.

The functional implication of IFN-γ signaling in cancer progression remains unclear[29, 30]. We showed that deletion of CCRL2/IFN-γ targets has a relatively negative impact on erythroleukemia/*TP53* MT AML cells survival. At the same time, exogenous IFN-γ decreases the survival of *TP53* MT AML blasts, suggesting a potentially different effect of exogenous and cell-intrinsic IFN-γ signaling activation. Significantly, CCRL2/IFN-γ targets upregulation was associated with the transformation of *TP53* MT pre-leukemic clones to multi-hit *TP53* MT AML and increased resistance to venetoclax consistent with previous findings[8, 10].

In conclusion, our results support that CCRL2 is overexpressed in AML with erythroid differentiation and *TP53* mutations inducing IFN-γ signaling response. Based on our studies, this is possibly a cell-intrinsic phenomenon related to p53 loss of function and independent of the bone marrow microenvironment. Activation of this pathway is associated with the selection of *TP53* MT pre-leukemic clones and resistance to venetoclax, supporting further investigation of CCRL2/IFN-γ signaling as a potential therapeutic target for AML with erythroid differentiation and *TP53* mutations.

## Materials and Methods

### Publicly available databases

Details on sample selection and data analysis are added in the *Online Supplementary Methods*.

### Patients and sample processing

Bone marrow samples were collected from bone marrow aspirations of patients with *TP53* mutations (at least 50% VAF) and complex karyotype. Normal marrow was collected as excess material after harvesting normal donors for allogenic bone marrow transplantation. All specimens were obtained by the Johns Hopkins Kimmel Cancer Center Specimen Accessioning Core. In accordance with the Declaration of Helsinki and under a research protocol approved by the Johns Hopkins Institutional Review Board, informed consent was procured from all donors before specimen collection. Isolation of CD34+ cell subsets was performed using the CD34 MicroBead kit (Miltenyi Biotec) as before[12, 13].

### Cell lines and reagents

Human TF-1, K562, SET2, HEL, and MV-411 cell lines were purchased from American Type Culture Collection. F36P cells were purchased from Leibniz Institute DSMZ. UKE-1 cells purchased from Coriell Institute. TF-1 cells and F36P cells were cultured in RPMI 1640 (Thermo Fischer Scientific, Waltham, MA) with 10% and 20% fetal bovine serum (FBS) (MilliporeSigma, Burlington, MA) respectively with the addition of GM-CSF (2 ng/ml and 20 ng/ml respectively; PeproTech). K562 and UKE-1 cells were cultured in RPMI 1640 with 10% FBS. SET2 and HEL cells were cultured in RPMI 1640 with 10% FBS. MV4-11 cells were cultured in IMDM (Thermo Fischer Scientific, Waltham, MA) with 10% FBS. All the cell lines were cultured with 2mM L-glutamine, penicillin (100 U/ml) and streptomycin (100µg/ml) at 37 in 5% CO2. Doxycycline was purchased from Sigma Aldrich (D9891) and was diluted in PBS (Thermo Fischer Scientific, Waltham, MA).

### Flow cytometry analysis

Cells from healthy controls and patients with AEL and AML cell lines were stained with a fluorescently conjugated antibody, PE-conjugated anti-human CCRL2 (BioLegend, #358303) and PE/Cy-7-conjugated anti-mouse CD45 (BioLegend #103113), BV510-conjugated anti-human CD45 (BioLegend, #103137). For the assessment of apoptosis, cells were stained with 7AAD (BioLegend, #420403). Gating was based on an unstained control. Following staining, analysis was performed using BD LSR II (BD Biosciences). Mean fluorescence intensity (MFI) was measured for each marker using FlowJo analysis software version 10.0.8(FlowJo, Ashland, CO, USA).

### CCRL2 and TP53 knockout

Lentiviral vectors expressing CCRL2-targeting sgRNA (pLV [CRISPR]-hCas9:T2A: Puro-U 6>hCCRL2[gRNA#162], pLV [CRISPR]-hCas9:T2A: Puro-U6>hCCRL2 [gRNA#177]) or empty pLKO.1-puro lentiviral vector (pLV [CRISPR]-hCas9/Puro-U6>Scramble_gRNA1) were transfected into 293T cells using Lipofectamine 2000 (Thermo Fischer Scientific) for lentiviral supernatant production. TF-1, F36P, K562, SET2, HEL and MV-411 cells were incubated with the viral supernatant and polybrene (8µg/ml; MilliporeSigma) for transduction. 48 hours later, cells were treated with puromycin (2µg/ml for TF-1 and K562, 1.5 µg/ml for F36P, 0.5 µg/ml for SET2 and HEL and 0.75 µg/ml for MV4-11) for 3-4 days for resistant cells selection.

Similarly, lentiviral vectors expressing TP53-targeting sgRNA (or empty were transfected into 293T cells as detailed above. UKE-1 cells were incubated with the viral supernatant and polybrene (8µg/ml; MilliporeSigma) for transduction. 48 hours later, cells were treated with Blasticidin (10 µg/ml) for 3-4 days for resistant cells selection.

### Colony formation assay

Clonogenic assays were conducted as previously detailed [12, 13]. Cells were counted and resuspended at a density of 3000 cells/ml in methylcellulose-based media. Following around two weeks of incubation at 37°C in 5% CO2, counting of colony forming units was performed under bright-field microscopy. A colony was defined as a cell aggregate of >50 cells.

### TF-1 and SET2 xenograft mice

After transduction of TF-1 and SET2 cells with sgRNAs, and selection of resistant cells as above, resistant cells were transduced with a GFP/Luciferase+ retroviral vector as before [12, 13]. Afterward, GFP+ cells sorted, and injected intravenously to 8-10-week-old NOD.Cg-*Prkdc^scid^ Il2rg^tm1Wjl^*/SzJ (NSG) female mice (10^6^ cells per mouse). Using IVIS spectrum in vivo imaging system, bioluminescence signal was measured. At day 65 for TF-1 and 60 for SET2, all remaining mice were euthanized, and the percentage of human CD45+ cells was assessed in bone marrow by flow cytometry.

### Mass spectrometry phosphoproteomics analysis

Protein extracts were buffer exchanged using SP3 paramagnetic beads (GE Healthcare)[43]. Briefly, protein samples (20 ug) were brought up to 100 uL with 10 mM TEAB + 1% SDS and disulfide bonds reduced with 10 uL of 50 mM dithiothreitol for 1 hour at 60C. Samples were cooled to RT and pH adjusted to ∼7.5, followed by alkylation with 10 uL of 100 mM iodoacetamide in the dark at RT for 15 minutes. Next, 100 ug (2 uL of 50 ug/uL) SP3 beads were added to the samples, followed by 120 uL 100% ethanol. Samples were incubated at RT with shaking for 5 minutes. Following protein binding, beads were washed with 180 uL 80% ethanol three times. Proteins were digested on-bead with 2ng trypsin (Pierce) in 100uL 25mM TEAB buffer at 37C overnight. Resulting peptides were separated from the beads using a magnetic tube holder. Supernatants containing peptides were acidify and desalted on u-HLB Oasis plates. Peptides were eluted with 60% acetonitrile/0.1%TFA and dried using vacuum centrifugation.

Each of the 12 dried peptide samples were labeled with one of the unique TMTpro 16-plex reagents (Thermo Fisher, Lot WK338750) according to the manufacturer’s instructions. All 12 TMT labeled peptide samples were combined and dried by vacuum centrifugation.

The combined TMT-labeled peptides were re-constituted in 100 µL 200mM TEAB buffer and filtered through Pierce Detergent removal columns (Fisher Scientific PN 87777) to remove excess TMT label. Peptides in the flow through were diluted to 2 mL in 10 mM TEAB in water and loaded on a XBridge C18 Guard Column (5 µm, 2.1 x 10 mm, Waters) at 250 µL/min for 8 min prior to fractionation on a XBridge C18 Column (5 µm, 2.1 x 100 mm column (Waters) using a 0 to 90% acetonitrile in 10 mM TEAB gradient over 85 min at 250 µL/min on an Agilent 1200 series capillary HPLC with a micro-fraction collector. Eighty-four 250 ul fractions were collected and concatenated into 24 fractions[44]. From each fraction, 10% was analysis for global quantitative proteomic comparison and normalization. The remaining 90% was combined into 12 fractions for phosphopeptide enrichment by binding to titanium dioxide (TiO2)[45].

TMT labeled peptides before and after phosphopeptide enrichment were analyzed by nanoflow reverse phase chromatography coupled with tandem mass spectrometry (nLCMS/MS) on an Orbitrap-Fusion Lumos mass spectrometer (Thermo Fisher Scientific) interfaced with an EasyLC1000 UPLC. Peptides will be separated on a 15 cm × 75 μm i.d. self-packed fused silica columns with ProntoSIL-120-5-C18 H column 3 µm, 120 Å (BISCHOFF) using an 2-90% acetonitrile gradient over 85 minutes in 0.1% formic acid at 300 nl per min and electrosprayed through a 1 µm emitter tip (New Objective) at 2500 V. Survey scans (MS) of precursor ions were acquired with a 2 second cycle time from 375-1500 m/z at 120,000 resolution at 200 m/z with automatic gain control (AGC) at 4e5 and a 50 ms maximum injection time. Top 15 precursor ions were individually isolated within 0.7 m/z by data dependent monitoring and 15s dynamic exclusion and fragmented using an HCD activation collision energy 39. Fragmentation spectra (MS/MS) were acquired using a 1e5 AGC and 118 ms maximum injection time (IT) at 50,000 resolution.

Fragmentation spectra were processed by Proteome Discoverer (v2.4, ThermoFisher Scientific) and searched with Mascot v.2.8.0 (Matrix Science, London, UK) against RefSeq2021_204 database. Search criteria included trypsin enzyme, two missed cleavage, 5 ppm precursor mass tolerance, 0.01 Da fragment mass tolerance, with TMTpro on N-terminus and carbamidomethylation on C as fixed modifications and TMTpro on K, phosphorylation on S, T or Y, oxidation on M, deamidation on N or Q as variable modifications. Peptide identifications from the Mascot searches were processed within PD2.4 using Percolator at a 5% False Discovery Rate confidence threshold, based on an auto-concatenated decoy database search. Peptide spectral matches (PSMs) were filtered for Isolation Interference <25%. Relative protein abundances of identified proteins were determined in PD2.4 from the normalized median ratio of TMT reporter ions from the top 30 most abundant proteins identified. ANOVA method was used to calculate the p-values of mean protein ratios for the biological replicates set up using a non-nested (or unpaired) design. Z-score transformation of normalized protein abundances from a quantitative proteomics analysis using isobaric mass tags was applied before performing the hierarchical clustering based on Euclidean distance and complete (furthest neighbors) linkage. The horizontal dendogram shows the proteins in samples that clustered together. Technical variation in ratios from our mass spectrometry analysis is less than 10% [46].

### Pathway Enrichment Analysis

Mass spectrometry experimental data were processed on the Proteome Discoverer platform, as described: abundance values were grouped and mapped to 7,173 unique proteins, then were imported into Partek Genomics Suite 6.6 (Partek Inc. Saint Louis MO) for further analysis and subsequent comparison of CCRL2 knockout.

These raw grouped values were then transformed to log2 notation and the wild-type and knock-down values were then quantile normalized to reduce experimental noise across the three biological replicates each. A t-test analysis was then run to compare the two, WT and KO, biological classes by relative abundance, expressed as fold change, and statistical significance, expressed as p-value. Proteins that demonstrated >2SD absolute value fold change were judged to be differentially abundant, and all these results were exported to the Ingenuity Pathway Analysis (QIAGEN Inc.) platform for functional analysis.

Ingenuity Pathway Analysis, IPA, evaluated its known pathways to determines those that are enriched for proteins that demonstrated this >2SD fold change, by comparing each pathway’s number of proteins that do or do-not not show this fold change. For each specific pathway IPA returned a Fisher’s exact test of whether an enrichment exists, and a Z-score whether the pathway is likely inhibited or activated.

### Western Blotting

Protein extraction was performed using M-PER^TM^ Mammalian Protein Extraction Reagent (#78501). Antibodies against P-STAT1 (Tyr^701^) (#9167), P-STAT1 (Ser^727^) (#9177), STAT1(#9172), and ßactin (#4970) were all purchased from Cell Signaling Technology.

### Co-immunoprecipitation

Cell lysates from TF-1, F36P, SET2 cells and doxy-inducible CCRL2 TF-1 cells were incubated overnight with Sepharose bead conjugate JAK2 monoclonal antibody (Cell Signaling Technology, #4089) or Sepharose bead conjugate isotype control (Cell Signaling Technology, #3423) as previously described[12]. Beads were then washed extensively and boiled with 30 µl of loading buffer.

### Single cell RNA sequencing analysis

Publicly available single cell RNA sequencing data from Kuusanmaki et al. was preprocessed as previously described [27]. Gene expression was visualized using Seurat (v5.1.0) and scCustomize (v2.0.1) in R (v4.3.1).

Another publicly available single cell RNA sequencing data from van Galen et al. [28]. Gene expression was visualized using Seurat (v5.1.0) (PMID 29608179) and in R (v4.3.1). Briefly, the single cell data across samples were merged. The UMI matrix was then normalized for library size, log transformed and scaled. Principal Component Analysis (PCA) was performed followed by Louvain nearest neighbor clustering using 30 PCs. UMAP was applied to visualize the cell distribution in 2D space. A resolution of 0.5 was used for clustering to identify different cell populations. Cluster identity was identified using canonical marker genes. Differential expression (DE) analysis in individual clusters across groups of samples was performed using a pseudobulk approach. Specifically, counts across cells for each sample were aggregated using the AggregateExpression function. Surrogate variable analysis (SVA) was applied to identify latent confounders (PMID 22257669) and differential expression was estimated using DESeq2 v1.44.0 (PMID 25516281) with surrogate variables as covariates. FDR < 0.05 was set as the significance threshold.

### Bulk RNA sequencing

We followed the DESeq2 pipeline (PMID 25516281) to normalize the gene expression results using a size factor that accounts for library size and gene size. This was followed by a variance stabilizing transformation as implemented in DESeq2, the output of which was used to perform principal components analysis (PCA). Surrogate variable analysis (SVA) was employed to control for unknown confounders and batch effects (PMID 22257669) while preserving the biological differences between groups. Differential expression (DE) was tested based on a Wald test with a two-tailed alternative hypothesis and a corresponding p-value was generated and adjusted for multiple testing using false discovery rate (FDR). Genes were considered significant if their FDR was < 0.05. The gene set enrichment analysis (GSEA) package was used to identify pathways enriched in our DE results (PMID 16199517).

### Quantitative real-time PCR

Quantitative real-time polymerase chain reaction (PCR) was performed as previously described[47]. Total RNA was extracted using the Monarch Total RNA miniprep kit(T2010S) and complementary DNA was synthesized using QuantiTect Rev. Transcription Kit (#205311, Qiagen, Valencia, CA). Quantitative real-time PCR was conducted by using sequence-specific primers. The Radiant SYBR Green Lo-ROX qPCR kit (Alkali Scientific, Fort Lauderdale, FL) and CFX96 real-time PCR detection system (Bio-Rad) were utilized. Normalization of RNA expression was based on GAPDH expression.

### Nuclear and cytoplasmic Fractionation

Nuclear/cytoplasmic fractionation was performed using the NE-PER^TM^ Nuclear and Cytoplasmic Extraction Kit (ThermoFischer Scientific #78833) as previously described[48].

### Statistical Analysis

Statistical analysis was performed by using GraphPad Prism (GraphPad Software, La Jolla, CA). Mann-Whitney test was performed to assess statistical significance when comparing two groups. One-way analysis of variance (ANOVA) was performed for the comparisons of three or more groups. Dunnett’s test was used for multiple comparisons. Standard deviation was used to assess centrality and dispersion.

## List of Supplementary Materials

**-Supplementary Materials and Methods**

**-Fig S1**

**-Tables S1**

## Author contributions

N.S.N, S.P. T.C. and T.K conceived and designed the study and wrote the manuscript. N.S.N, T.C., B.P., C.E. performed the flow cytometry analysis. N.S.N, Y.H., X.Z., F.B. and T.K. performed the CRISPR-Cas9 deletion experiments. N.S.N, T.C., and Y.H. performed the Western blot analysis. N.S.N, Y.H. and X.Z. performed the quantitative real time PCR. N.S.N., S.P., T.C., L.S.R. and L.G. analyzed publicly available bulk RNA sequencing data, J.R., P.T.S., I.G., L.L. and M.A. analyzed publicly available single-cell RNA sequencing data, T.R.B. and R.N.C. performed the phospho-proteomics analysis, C.T. performed the ingenuity pathway analysis, P.T.S and M.A. performed the gene-set-enrichment analysis, N.S.N., B.P. and T.K. processed the primary samples, N.S.N., T.C., I.S., P.T. S.K., and T.K. performed the xenograft studies. N.S.N., B.P., B.C.P. and T.K. processed the primary samples. N.S.N, B.P., C.E. and T.K. performed the clonogenic assays, A.A. and B.P. performed ELISA, P.F., A.J.A., M.J.L, A.E.D., R.J.J. identified patients’ samples and provided clinical information, C.L. and R.X. provided pathologic assessment for primary samples, N.S.N., T.K., M.J.L., S.K., L.S.R., A.E.D., R.J.J., F.B., S.P. interpreted the data and edited the manuscript.

## Data and materials availability

All data are available in the main text or the supplementary materials.

## REFERENCES

1. Alaggio, R., et al., The 5th edition of the World Health Organization Classification of Haematolymphoid Tumours: Lymphoid Neoplasms. Leukemia, 2022. 36(7): p. 1720–1748.

2. Almeida, A.M., et al., Clinical Outcomes of 217 Patients with Acute Erythroleukemia According to Treatment Type and Line: A Retrospective Multinational Study. Int J Mol Sci, 2017. 18(4).

3. Daver, N.G., et al., TP53-Mutated Myelodysplastic Syndrome and Acute Myeloid Leukemia: Biology, Current Therapy, and Future Directions. Cancer Discov, 2022. 12(11): p. 2516–2529.

4. Fang, H., et al., Pure erythroid leukemia is characterized by biallelic TP53 inactivation and abnormal p53 expression patterns in de novo and secondary cases. Haematologica, 2022. 107(9): p. 2232–2237.

5. Reichard, K.K., et al., Pure (acute) erythroid leukemia: morphology, immunophenotype, cytogenetics, mutations, treatment details, and survival data among 41 Mayo Clinic cases. Blood Cancer J, 2022. 12(11): p. 147.

6. Wong, T.N., et al., Role of TP53 mutations in the origin and evolution of therapy-related acute myeloid leukaemia. Nature, 2015. 518(7540): p. 552-555.

7. Sinanidis, I., et al., Favorable outcomes in MDS and oligoblastic AML-MR after reduced-intensity conditioning allogeneic bone marrow transplantation with post-transplantation cyclophosphamide. Bone Marrow Transplant, 2024. 59(8): p. 1178–1180.

8. Rodriguez-Meira, A., et al., Single-cell multi-omics identifies chronic inflammation as a driver of TP53-mutant leukemic evolution. Nat Genet, 2023. 55(9): p. 1531–1541.

9. Vadakekolathu, J., et al., TP53 abnormalities correlate with immune infiltration and associate with response to flotetuzumab immunotherapy in AML. Blood Adv, 2020. 4(20): p. 5011–5024.

10. Wang, B., et al., Comprehensive characterization of IFNγ signaling in acute myeloid leukemia reveals prognostic and therapeutic strategies. Nat Commun, 2024. 15(1): p. 1821.

11. Schioppa, T., et al., Molecular Basis for CCRL2 Regulation of Leukocyte Migration. Front Cell Dev Biol, 2020. 8: p. 615031.

12. Karantanos, T., et al., The role of the atypical chemokine receptor CCRL2 in myelodysplastic syndrome and secondary acute myeloid leukemia. Sci Adv, 2022. 8(7): p. eabl8952.

13. Karantanos, T., et al., CCRL2 aZects the sensitivity of myelodysplastic syndrome and secondary acute myeloid leukemia cells to azacitidine. Haematologica, 2023. 108(7): p. 1886–1899.

14. Bagger, F.O., et al., BloodSpot: a database of gene expression profiles and transcriptional programs for healthy and malignant haematopoiesis. Nucleic Acids Res, 2016. 44(D1): p. D917–24.

15. Tyner, J.W., et al., Functional genomic landscape of acute myeloid leukaemia. Nature, 2018. 562(7728): p. 526-531.

16. Boehm, J.S., et al., Cancer research needs a better map. Nature, 2021. 589(7843): p. 514-516.

17. Khoury, J.D., et al., The 5th edition of the World Health Organization Classification of Haematolymphoid Tumours: Myeloid and Histiocytic/Dendritic Neoplasms. Leukemia, 2022. 36(7): p. 1703–1719.

18. Li, B., et al., BMP2/SMAD pathway activation in JAK2/p53-mutant megakaryocyte/erythroid progenitors promotes leukemic transformation. Blood, 2022. 139(25): p. 3630–3646.

19. Ma, F., et al., Retinoid X receptor α attenuates host antiviral response by suppressing type I interferon. Nat Commun, 2014. 5: p. 5494.

20. Zelcer, N. and P. Tontonoz, Liver X receptors as integrators of metabolic and inflammatory signaling. J Clin Invest, 2006. 116(3): p. 607–14.

21. Subramanian, A., et al., Gene set enrichment analysis: a knowledge-based approach for interpreting genome-wide expression profiles. Proc Natl Acad Sci U S A, 2005. 102(43): p. 15545–50.

22. Mao, X., et al., Structural bases of unphosphorylated STAT1 association and receptor binding. Mol Cell, 2005. 17(6): p. 761–71.

23. Ivashkiv, L.B. and L.T. Donlin, Regulation of type I interferon responses. Nat Rev Immunol, 2014. 14(1): p. 36–49.

24. Nguyen, H., et al., Roles of phosphatidylinositol 3-kinase in interferon-gamma-dependent phosphorylation of STAT1 on serine 727 and activation of gene expression. J Biol Chem, 2001. 276(36): p. 33361–8.

25. Kovarik, P., et al., Stress-induced phosphorylation of STAT1 at Ser727 requires p38 mitogen-activated protein kinase whereas IFN-gamma uses a diZerent signaling pathway. Proc Natl Acad Sci U S A, 1999. 96(24): p. 13956–61.

26. Quelle, F.W., et al., Phosphorylation and activation of the DNA binding activity of purified Stat1 by the Janus protein-tyrosine kinases and the epidermal growth factor receptor. J Biol Chem, 1995. 270(35): p. 20775–80.

27. Kuusanmäki, H., et al., Erythroid/megakaryocytic diZerentiation confers BCL-XL dependency and venetoclax resistance in acute myeloid leukemia. Blood, 2023. 141(13): p. 1610–1625.

28. van Galen, P., et al., Single-Cell RNA-Seq Reveals AML Hierarchies Relevant to Disease Progression and Immunity. Cell, 2019. 176(6): p. 1265–1281.e24.

29. Mandai, M., et al., Dual Faces of IFNγ in Cancer Progression: A Role of PD-L1 Induction in the Determination of Pro- and Antitumor Immunity. Clin Cancer Res, 2016. 22(10): p. 2329–34.

30. Beziaud, L., et al., IFNγ-induced stem-like state of cancer cells as a driver of metastatic progression following immunotherapy. Cell Stem Cell, 2023. 30(6): p. 818–831.e6.

31. Iacobucci, I., et al., Genomic subtyping and therapeutic targeting of acute erythroleukemia. Nat Genet, 2019. 51(4): p. 694–704.

32. Iacobucci, I., et al., Modeling and targeting of erythroleukemia by hematopoietic genome editing. Blood, 2021. 137(12): p. 1628–1640.

33. Takeda, J., et al., Amplified EPOR/JAK2 Genes Define a Unique Subtype of Acute Erythroid Leukemia. Blood Cancer Discov, 2022. 3(5): p. 410–427.

34. Gera, K., et al., Survival after Pure (Acute) Erythroid Leukemia in the United States: A SEER-Based Study. Cancers (Basel), 2023. 15(15).

35. Weinberg, O.K. and D.A. Arber, Erythroleukemia: an Update. Curr Oncol Rep, 2021. 23(6): p. 69.

36. Boddu, P., et al., Erythroleukemia-historical perspectives and recent advances in diagnosis and management. Blood Rev, 2018. 32(2): p. 96–105.

37. Takeda, J., [Molecular pathogenesis and therapeutic targets in acute erythroid leukemia]. Rinsho Ketsueki, 2022. 63(2): p. 121–133.

38. Del Prete, A., et al., The atypical receptor CCRL2 is required for CXCR2-dependent neutrophil recruitment and tissue damage. Blood, 2017. 130(10): p. 1223–1234.

39. Marki, A. and K. Ley, Leaking chemokines confuse neutrophils. J Clin Invest, 2020. 130(5): p. 2177–2179.

40. Stengel, A., et al., Interplay of TP53 allelic state, blast count, and complex karyotype on survival of patients with AML and MDS. Blood Adv, 2023. 7(18): p. 5540–5548.

41. Ramana, C.V., et al., Stat1-dependent and -independent pathways in IFN-gamma-dependent signaling. Trends Immunol, 2002. 23(2): p. 96–101.

42. Li, L., et al., TP53 Mutations within T Cells Induce T-Cell Exhaustion and Functional Impairment in TP53 Mutant AML. Blood, 2024. 144(Supplement 1): p. 330–330.

43. Hughes, C.S., et al., Single-pot, solid-phase-enhanced sample preparation for proteomics experiments. Nat Protoc, 2019. 14(1): p. 68–85.

44. Wang, Y., et al., Reversed-phase chromatography with multiple fraction concatenation strategy for proteome profiling of human MCF10A cells. Proteomics, 2011. 11(10): p. 2019–26.

45. Larsen, M.R., et al., Highly selective enrichment of phosphorylated peptides from peptide mixtures using titanium dioxide microcolumns. Mol Cell Proteomics, 2005. 4(7): p. 873–86.

46. Herbrich, S.M., et al., Statistical inference from multiple iTRAQ experiments without using common reference standards. J Proteome Res, 2013. 12(2): p. 594–604.

47. Chang, Y.T., et al., Role of CYP3A4 in bone marrow microenvironment-mediated protection of FLT3/ITD AML from tyrosine kinase inhibitors. Blood Adv, 2019. 3(6): p. 908–916.

48. Hung, S.C., et al., Angiogenic eZects of human multipotent stromal cell conditioned medium activate the PI3K-Akt pathway in hypoxic endothelial cells to inhibit apoptosis, increase survival, and stimulate angiogenesis. Stem Cells, 2007. 25(9): p. 2363–70.

