## Supplementary Figure Legends for "CCRL2 promotes the interferon-γ signaling response in myeloid neoplasms with erythroid differentiation and mutated *TP53*"

**Supplementary Fig. 1. (A)** Lollipop plot showing specific TP53 variants of the patients’ samples (**B)** Gating strategy for analysis of healthy donors’ samples based on CD34/CD71/CD235a expression. **(C)** Gating strategy for analysis of TP53 mutated MDS/AML and AEL samples based on SSC-A/CD45 to identify blasts which express high CD71/CCRL2 (for AEL patients) and CD34/CD71 expression to identify CD34+CD71+ cells (for AEL patients). **(D-E)** *TP53* MT MDS/AML patients with EP showed a trend toward higher CCRL2 expression in their blasts and CD34+ cells respectively compared to *TP53* MT MDS/AML without EP.

**Supplementary Fig. 2.** **(A)** Western blot confirming the TP53 knockouts (KO1 and KO2) in UKE-1 cells. **(B)** qPCR showing higher CCRL2 expression in *TP53* KO UKE-1 cells compared to *TP53* WT UKE-1 cells. **(C-H)** CCRL2 knockout (KO) with two different lentiviruses (sgCCRL2 1 and sgCCRL2 2) compared to scramble sgRNA (sgControl) was confirmed by flow cytometry in TF-1, F36P, K562, SET2 and HEL cells respectively. **(I)** CCRL2 KO showed no effect on the clonogenicity of the *TP53* WT monocytic MV4-11 cells.

**Supplementary Fig. 3. (A)** Protein-protein interaction network showing that CCRL2 positively regulates IFN-γ signaling and suppresses LXR/RXR activation. **(B)** Bulk RNA-seq was performed followed by gene set enrichment analysis (GSEA) using a compilation of pathways from MSigDB in CCRL2 WT or KO SET2 cells. Volcano plot portrays the expression of the differentially expressed genes. **(C)** Principal components analysis showing principal components 1 and 2 (PC1 and PC2) variances for SET2 WT and KO cells. **(D)** Heatmap shows that IFN-γ gene networks was found to be the top CCRL2-regulated pathway.

**Supplementary Fig. 4. (A)** CCRL2 KO decreased the nuclear translocation of STAT1 in TF-1 cells (C: cytoplasm, N: nucleus). **(B)** Treatment of TF-1 CCRL2 KO and WT cells with 10 ng/ml and 20 ng/ml of IFN-γ showed that CCRL2 KO had no effect on the upregulation of *IFIT3* expression as a response to exogenous IFN-γ. **(C)** Doxycycline (doxy)-inducible CCRL2 TF-1 cell model upon treatment with 10 or 100 ng/ml doxycycline induced CCRL2 expression and increased their growth at 2 and 4 days (p=0.008 for 10 ng/ml and p=0.003 for 100ng/ml).

**Supplementary Fig. 5. (A)** Comparison of the expression of *IFIT1*, *ISG15* and *IFIT3* between *TP53* WT and *TP53* MT AML samples was done by analyzing RNA sequencing data derived from Beat AML dataset. *IFIT1* was found to be significantly overexpressed in *TP53* MT AML samples compared to *TP53* WT ones (p=0.024). All three CCRL2/STAT1 target genes were upregulated in AML samples compared to healthy bone marrow mononuclear cells (p<0.001). **(B)** Analysis of single-cell RNA sequencing data from Kuusanmäki et al. showed that blasts with erythroid differentiation express higher levels of various CCRL2/IFN-γ targets including *IFIT1*, *IFIT2*, *IFIT3*, *IFI35*, *IFIH1* and *ISG15* compared to other AML differentiation clusters. **(C)** Analysis of single cell RNA sequencing data from van Galen et al. identified blasts from bone marrow aspirates of 16 AML patients including 3 individuals with *TP53* mutations by CD34 and c-KIT (CD117) expression and T-cells by CD3 expression. **(D)** Comparison of *IFN*-γ in T-cells of *TP53* WT and MT AML patients based on single cell RNA sequencing data from van Galen et al. demonstrated that patients with *TP53* MT AML expressed relatively lower levels of *IFN-* γ compared to the WT ones without difference reaching statistical significance.

**Supplementary Fig. 6. (A**) Sorting of CD3+CD45+ and CD3-CD45+ cells from 4 *TP53* mutated AML patients and 3 healthy bone marrow donors. Blasts by dim CD45 and low side scatter were also sorted from the *TP53* mutated AML patients. **(B)** Cells (50,000 cells/ml) were cultured for 72 hours and levels of IFN-γ were then measured by ELISA, showing that T-cells from *TP53* MT AML cells secreted relatively lower IFN-γ levels compared to healthy donors. **(C)** Analysis of publicly available single-cell RNA sequencing data showed that pre-leukemic *TP53* heterozygous clones from patients with myeloproliferative neoplasms (MPN) who transformed to multi-hit *TP53* mutated sAML have higher expression of *IFIT1* (p<0.0001), *IFIT3 (*p<0.0001) and *ISG15* (p<0.0001) compared to *TP53* heterozygous clones from patients who remained in chronic phase (CP-MPN). **(D)** Heatmap showing the unsupervised clustering of genes and patients, considering the scaled gene expression values. Both rows and columns of the heatmap were cut in two based on dendrogram height. Through this, we separated the patients in two clusters based on the expression pattern of the CCRL2/IFN-γ targets: Cluster 2 patients (n = 35) who presented a high expression of *IFIT2, IFIT1, IFIT3, STAT1, IFI6, IFIH1, ISG15, IRF7*, whereas Cluster 1 patients (n = 134) contained the rest of the patients.
