## Supplementary Materials and Methods for "CCRL2 promotes the interferon-γ signaling response in myeloid neoplasms with erythroid differentiation and mutated *TP53*"

**Publicly available databases**

Analysis of the mRNA levels of *CCRL2* and STAT1 target genes (*IFIT1, ISG15 and IFIT3*) in different subtypes of AML was based on RNA sequencing data derived from The Cancer Genome ATLAS (TCGA). Clinical and transcriptomic data in the form of RSEM values from the TCGA LAML cohort were downloaded via cBioPortal[43-45]. Only samples that had transcriptomic and FAB information available were included. Patients were grouped based on their FAB status in M6/7 and M1/2/3/4/5, the latter being labelled as "Other". A total of 171 patients from the TCGA were included. Of those, 5 (2.9%) had an FAB of M6/7. Normality of the distribution of the RSEM values was assessed using skewness and kurtosis assessment and histogram visualization. Differences between two non-normally distributed groups were evaluated using the Mann-Whitney-Wilcoxon rank sum test. A p-value under 0.05 was considered statistically significant. We calculated an overall CCRL2/IFN signaling score based on TCGA dataset using an 18 gene list: RSEM values for CCRL2, STAT1, IFIT1, ICAM1, CD44, IFIT2, PRKCD, IFIT3, IFI35, ISG15, GBP2, IFIH1, MAPK14, GBP4, IFI6, IRF3, IRF7, IFITM3 were min max scaled at the gene level. To obtain a score for every patient, we took the median of these scaled values at the patient level. Similarly, data was extracted from DepMap Portal (DepMap22Q2). TPM values for CCRL2, STAT1, IFIT1, ICAM1, CD44, IFIT2, PRKCD, IFIT3, IFI35, ISG15, GBP2, IFIH1, MAPK14, GBP4, IFI6, IRF3, IRF7, IFITM3 were min max scaled at the gene level. To obtain a score for every patient, we took the median of these scaled values at the cell line level. Using Beat AML dataset, data was extracted from <http://vizome.org/aml/expression_strat/>. Patients were included if DxAtSpecimenAcquisition was Acute Myeloid Leukaemia (AML) and related precursor neoplasms or Healthy, pooled CD34+. Patients were included if Included_2018_DNAseqAnalysis was y or Healthy; pooled CD34+ and if none of the 18 included genes had n/a in the Normalized_RPKM column. A total of 407 patients were included. Of those, 12 (2.9%) were healthy CD34+, 360 (88.5%) were TP53 WT AML and 35 (8.6%) were TP53 MUT AML.Values from the RPKM_normalized column for CCRL2, STAT1, IFIT1, ICAM1, CD44, IFIT2, PRKCD, IFIT3, IFI35, ISG15, GBP2, IFIH1, MAPK14, GBP4, IFI6, IRF3, IRF7, IFITM3 were min max scaled at the gene level. To obtain a score for every patient, we took the median of these scaled values at the patient level. For overall survival using TCGA dataset, the surv_cutpoint function from the survminer package was used to determine an optimal cutoff for the CCRL2/IFN signaling score when it comes to overall survival. A patient was considered to have a high CCRL2/IFN signaling score if this score was over 0.1332826. The optimal cutoff for the RSEM values were 127.4210 for CCRL2, 161.9048 for IFIT1, 132.8000 for IFIT3, and 213.1177 for ISG15.

To determine specific expression differences (*CCRL2*, *IFIT1*, *IFIT3* and *ISG15*) between different CP-MPNs genotypes that progressed to sAML (pre-TP53-sAML) and that did not (CP-TP53-MPN), we used a previously published dataset[8]. Normalized gene expression per cell were downloaded from GSE226340 while corresponding genotyping data was downloaded from <https://doi.org/10.5281/zenodo.8060602>,metadata_MPNAMLp53_with_index_genotype.revised.txt.Normality of the distribution was assessed using histogram visualization and kurtosis and skewness evaluation. Differences between two non-normally distributed groups were assessed using Mann-Whitney-Wilcoxon rank sum test. Similarly, RNA expression data from DepMap Portal was analyzed to compare gene expression between different AML cell lines and data derived from Beat AML database was used to compare the expression of CCRL2, and STAT1 targets across AML samples based on molecular profile.

For data on the effect of IFN on venetoclax resistance, we included 169 patients. If a patient presented multiple samples, we included the sample with the lower ID number. We focused on the expression of the following genes: CCRL2, STAT1, IFIT1, ICAM1, CD44, IFIT2, PRKCD, IFIT3, IFI35, ISG15, GBP2, IFIH1, MAPK14, GBP4, IFI6, IRF3, IRF7, IFITM3. The expression values were min max scaled at a gene level. The CCRL2 Signaling Score was defined at a patient level and was considered as the median scaled expression level of the included genes. Correlation between the CCRL2 Signaling Score and Venetoclax IC50 was assessed using Spearmans rho. Unsupervised clustering and its graphical representation was performed using the pheatmap package ([https://github.com/raivokolde/pheatmap](https://nam02.safelinks.protection.outlook.com/?url=https%3A%2F%2Fgithub.com%2Fraivokolde%2Fpheatmap&data=05%7C02%7Cnnaji1%40jh.edu%7C2c6309caca7441276f0108dd33ff5f85%7C9fa4f438b1e6473b803f86f8aedf0dec%7C0%7C0%7C638723894168670001%7CUnknown%7CTWFpbGZsb3d8eyJFbXB0eU1hcGkiOnRydWUsIlYiOiIwLjAuMDAwMCIsIlAiOiJXaW4zMiIsIkFOIjoiTWFpbCIsIldUIjoyfQ%3D%3D%7C0%7C%7C%7C&sdata=QAHZUsO2EfbZ%2BGGeeWJr0qq78PN0%2Fa1SzQasRb9EAKs%3D&reserved=0)), using ward.D2 clustering. Mann-Whitney-Wilcoxon rank sum test was used to determine the difference in Venetoclax IC50 between Cluster 1 and 2.
