## Supplementary Figures for "CCRL2 promotes the interferon-γ signaling response in myeloid neoplasms with erythroid differentiation and mutated *TP53*"

### Slide 1
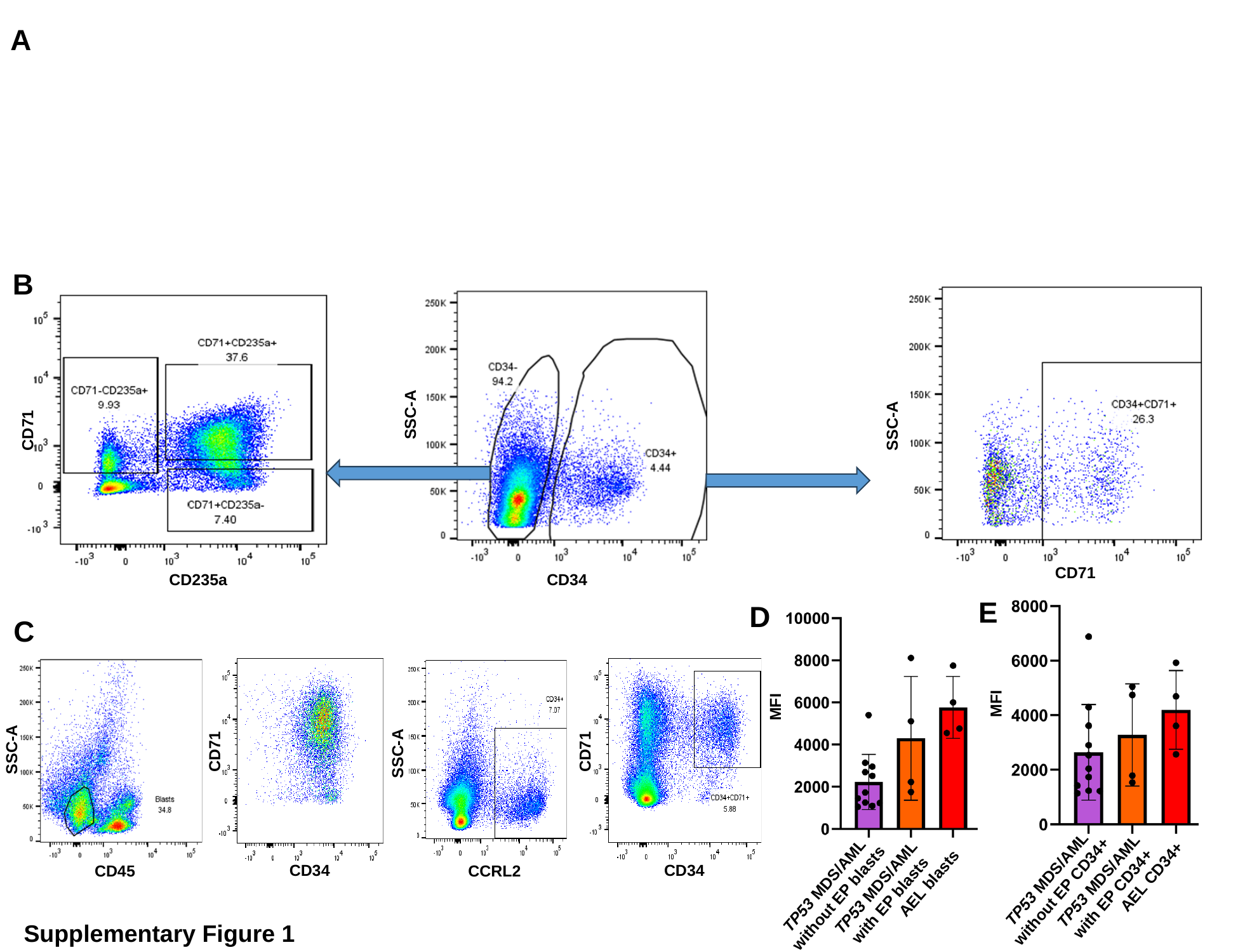

A
B
SSC-A
CD34
SSC-A
CD71
CD71
CD235a
E
D
C
SSC-A
SSC-A
CD71
CD71
MFI
MFI
CD34
CD34
CD45
CCRL2
TP53 MDS/AML without EP CD34+
AEL CD34+
TP53 MDS/AML with EP CD34+
TP53 MDS/AML without EP blasts
AEL blasts
TP53 MDS/AML with EP blasts
Supplementary Figure 1

### Slide 2
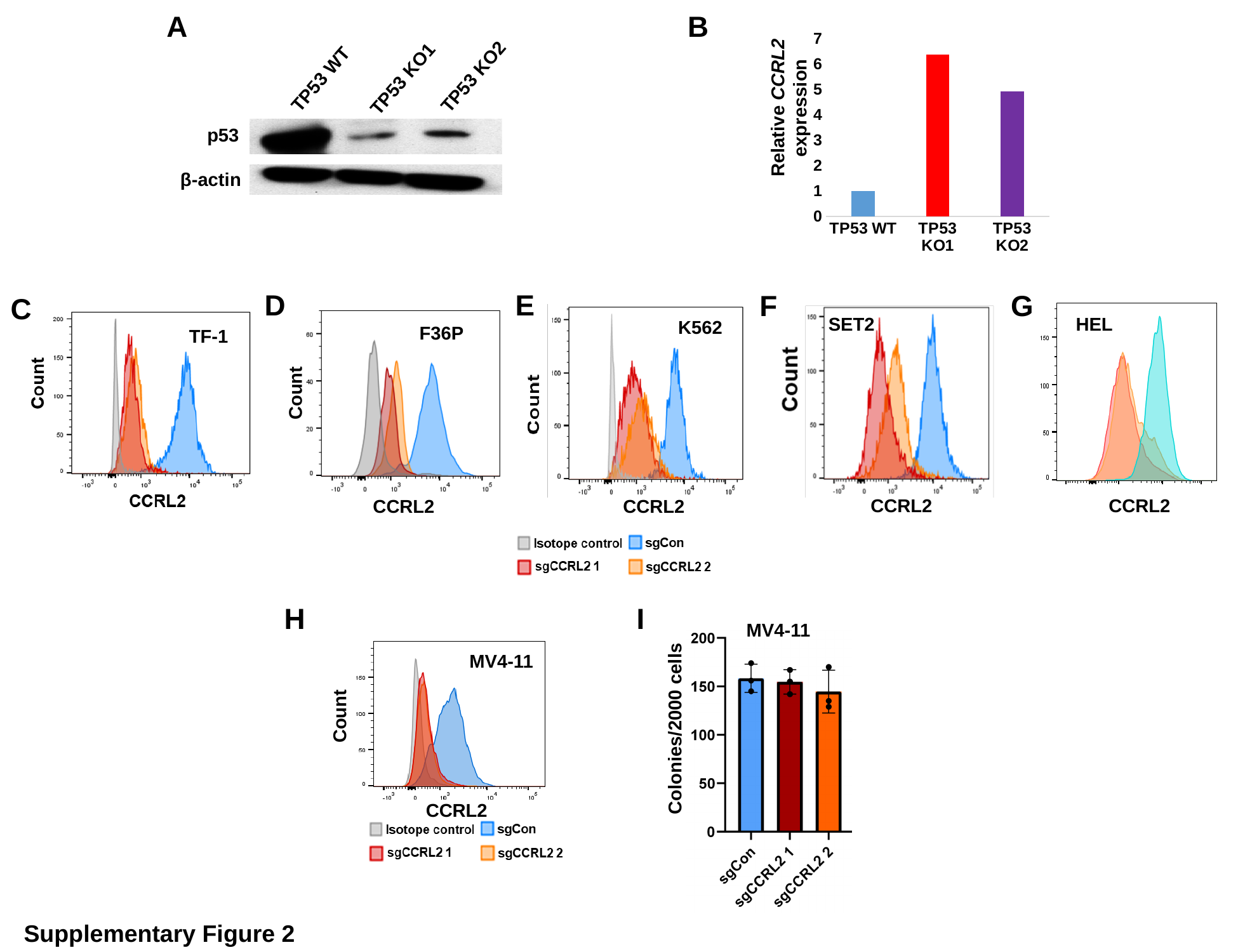

B
A
#### Chart
| Category | |
|---|---|
| TP53 WT | 1.0 |
| TP53 KO1 | 6.3790481677032025 |
| TP53 KO2 | 4.923545259119956 |TP53 WT
TP53 KO2
TP53 KO1
Relative CCRL2 expression
p53
β-actin
E
D
F
G
C
K562
HEL
SET2
Count
F36P
TF-1
CCRL2
CCRL2
CCRL2
CCRL2
H
I
MV4-11
Colonies/2000 cells
MV4-11
Count
CCRL2
Supplementary Figure 2

### Slide 3
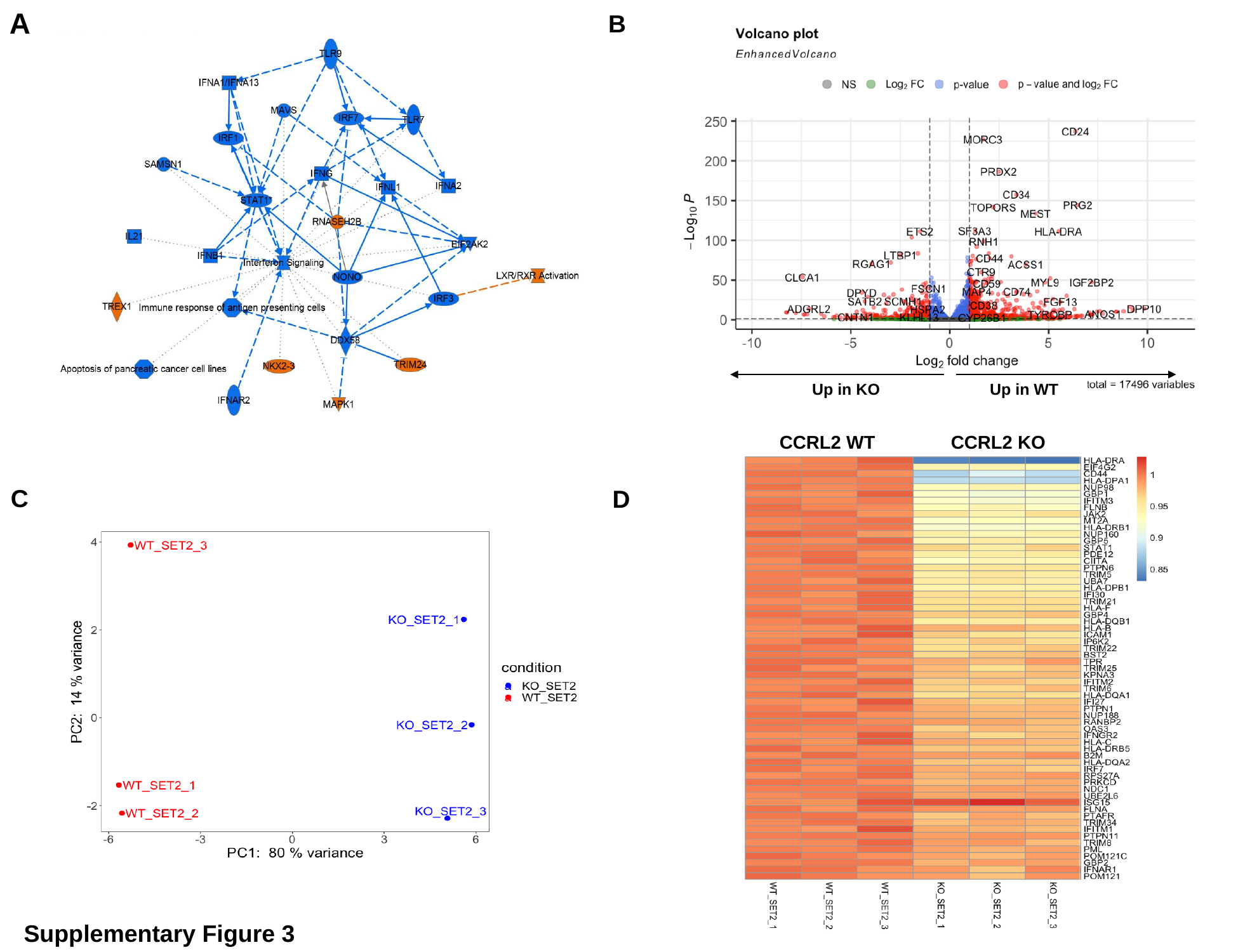

A
B
Up in WT
Up in KO
CCRL2 KO
CCRL2 WT
C
D
Supplementary Figure 3

### Slide 4
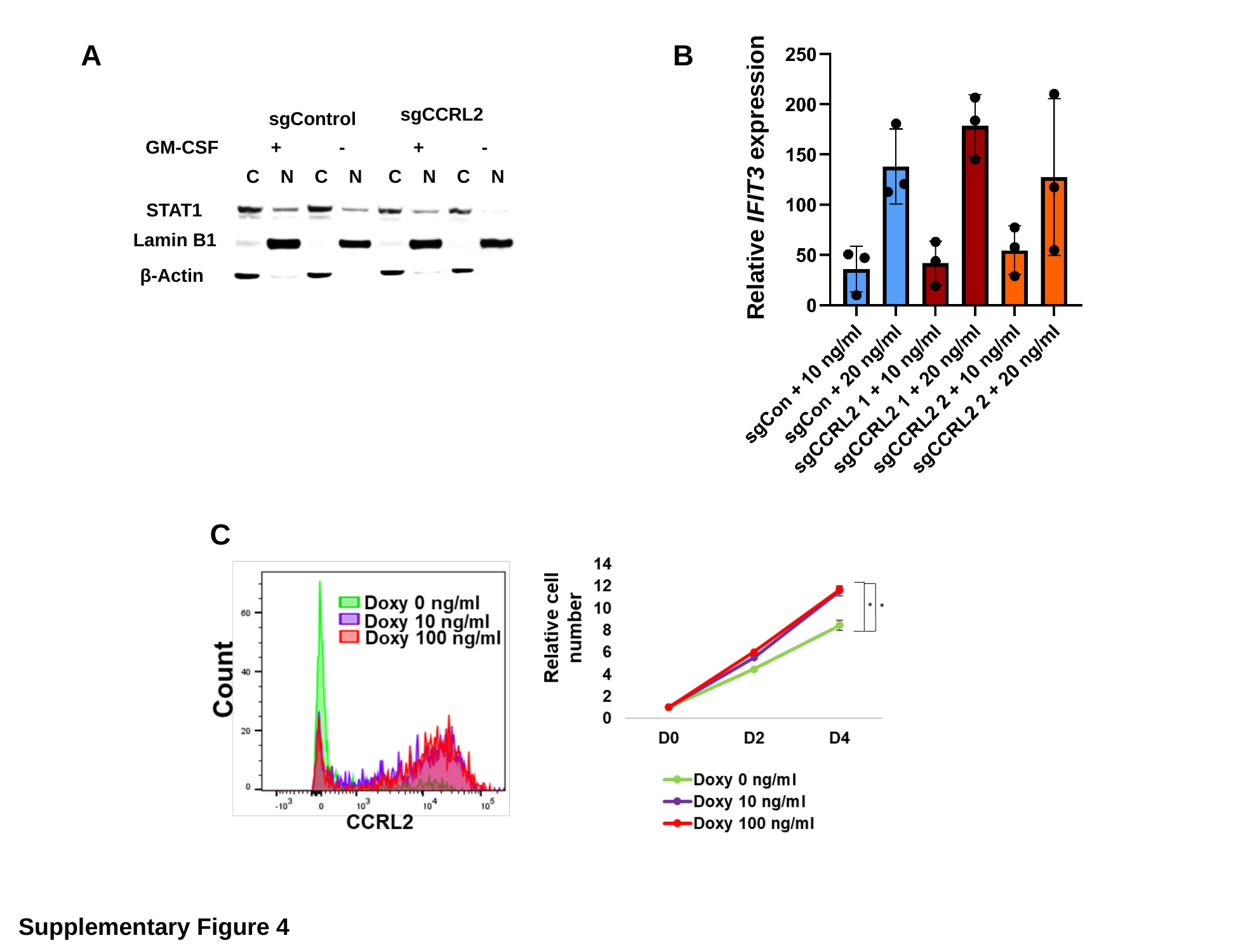

Relative IFIT3 expression
B
A
sgCCRL2
sgControl
GM-CSF + - + -
 C N C N C N C N
STAT1
Lamin B1
β-Actin
C
Supplementary Figure 4

### Slide 5
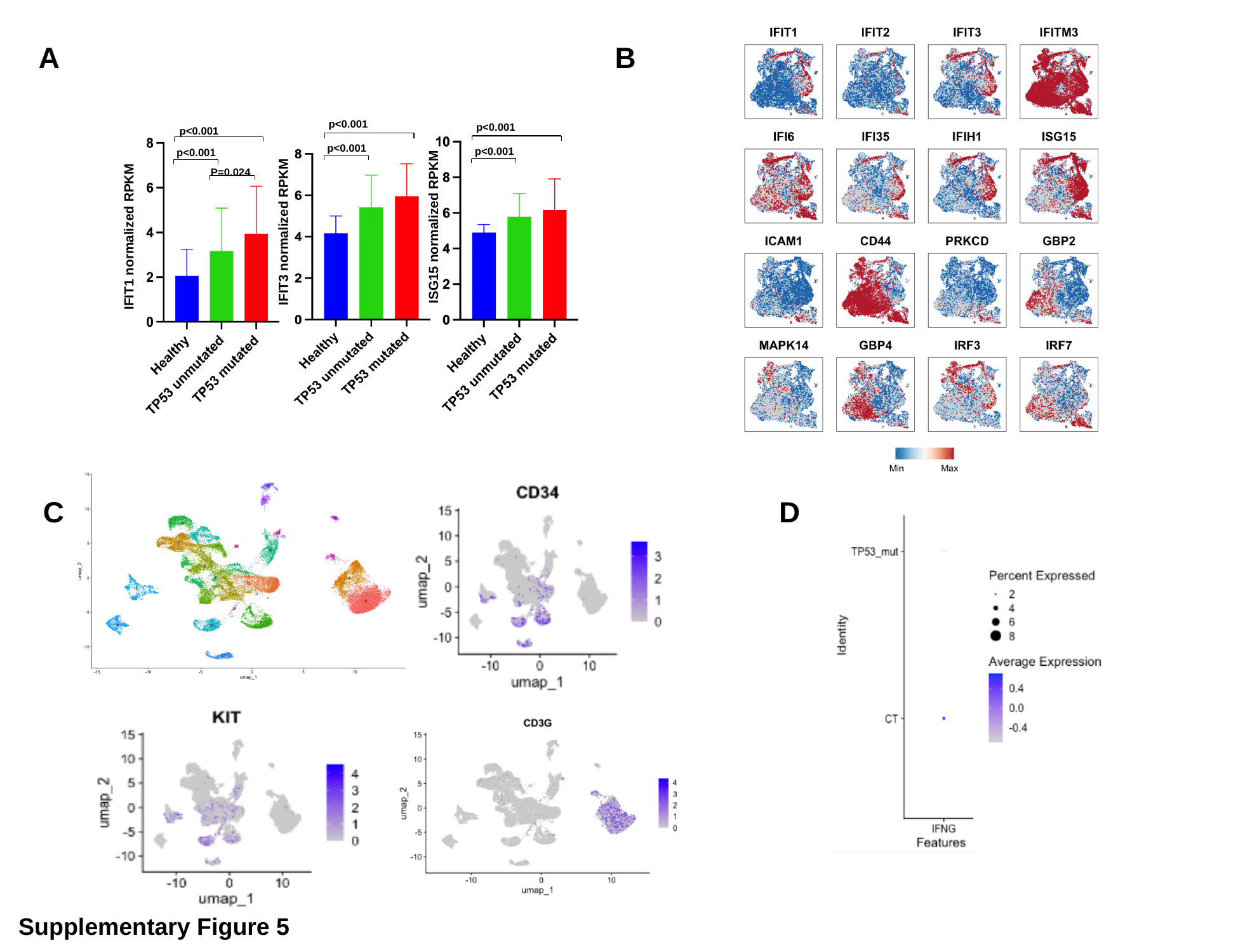

B
Α
p<0.001
p<0.001
p<0.001
p<0.001
p<0.001
p<0.001
P=0.024
ISG15 normalized RPKM
IFIT3 normalized RPKM
IFIT1 normalized RPKM
C
D
Supplementary Figure 5

### Slide 6
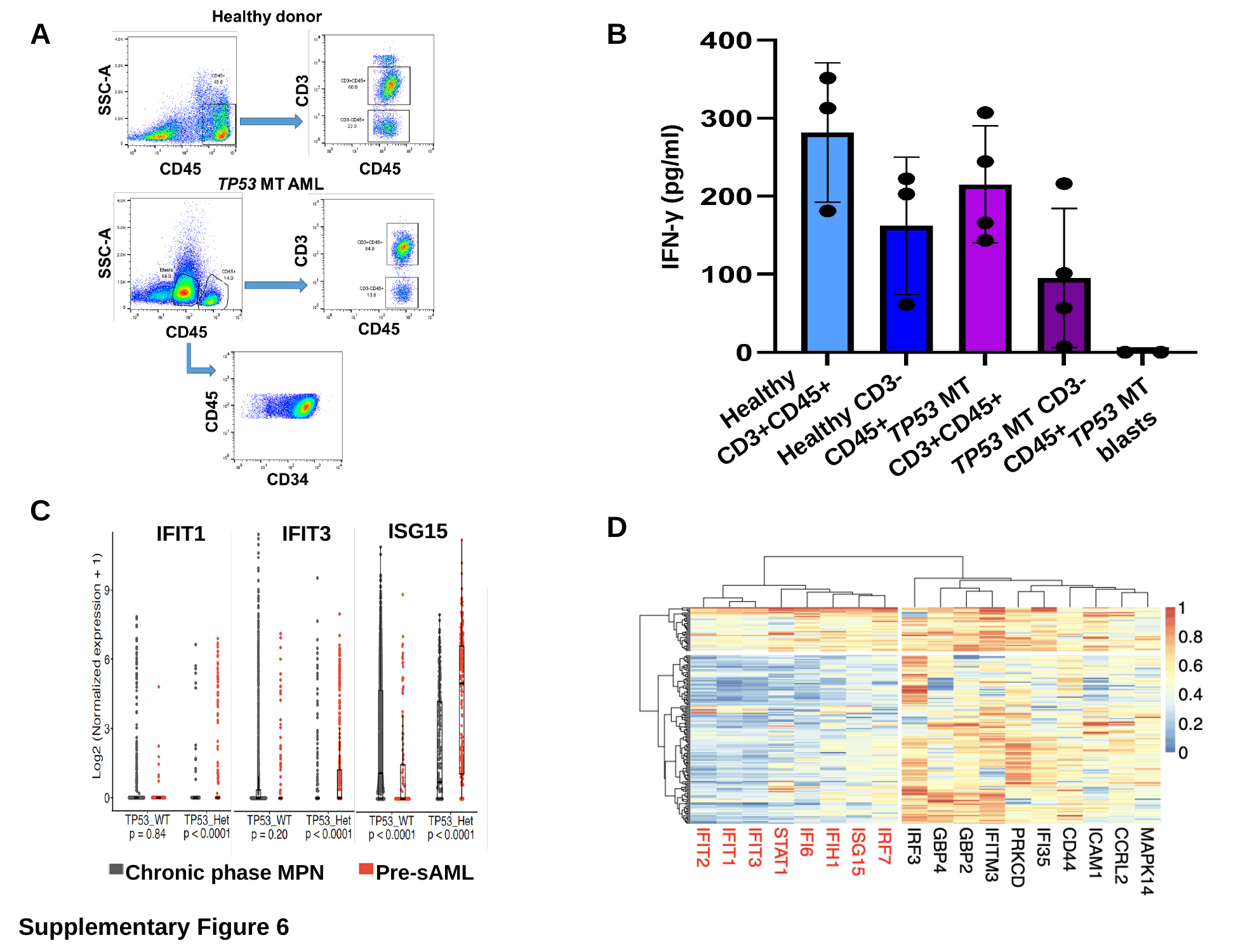

A
B
IFN-γ (pg/ml)
Healthy CD3+CD45+
TP53 MT CD3+CD45+
TP53 MT blasts
Healthy CD3-CD45+
TP53 MT CD3-CD45+
C
D
ISG15
IFIT1
IFIT3
Pre-sAML
Chronic phase MPN
Supplementary Figure 6
