## Supplementary Tables for "CCRL2 promotes the interferon-γ signaling response in myeloid neoplasms with erythroid differentiation and mutated *TP53*"

| **Characteristic** | ***TP53* mutated myeloid neoplasms (N=19)** | **Healthy donors (N=16)** |
| --- | --- | --- |
| Age | 65 (27 – 78) | 44 (30 – 56) |
| Gender  Females  Males | 4 (21%)  15 (79%) | 6 (37.5%)  10 (62.5%) |
| Diagnosis  *TP53* mutated MDS/AML  AEL | 15 (79%)  4 (21%) | N/A |
| Treatment-related | 6 (32%) | N/A |
| Blasts% | 25 (5-80) | N/A |
| Karyotype  Complex | 19 (100%) | N/A |

**Supplementary Table 1.** Characteristics of patients with *TP53* mutated myeloid neoplasms and healthy donors

| Variant(s) | VAF1 | VAF2 |
| --- | --- | --- |
| R158H | 79.9 |  |
| Y126H | 81.9 |  |
| D281N | 60.64 |  |
| H178P | 82.41 |  |
| Y220C, R158H | 24.5 | 26.7 |
| G245S | 67.5 |  |
| F270C | 89.9 |  |
| Y220C | 64.1 |  |
| V272M | 60.5 |  |
| R175H | 50.3 |  |
| c.994-1G>A | 85.24 |  |
| D281N | 97.54 |  |
| C275Y | 52.56 |  |
| S303fs | 86.5 |  |
| V272M | 69.14 |  |
| R196P | 50.2 |  |
| R273H, E285fs | 15.98 | 14.4 |
| H179R | 79.3 |  |
| E286G, R273H | 37.83 | 36.07 |

**Supplementary Table 2.** Variants and variant allele frequencies of patients with *TP53* mutated myeloid neoplasms.
